# Sex-specific effects of exercise on motor coordination and extended basal ganglia physiology

**DOI:** 10.64898/2026.04.24.720719

**Authors:** Valerie J. Lewitus, Lindsey A. Russ, Chelsea B. Scott, Olivia M. Kruszewski, Giorgi Shautidze, Zachary A. Colon, Shirley Q. Zhang, Joel A. Walker, Asia Fernandez, Megan R. Croom, Valeria G. Aleman, Rebekah C. Evans

## Abstract

Exercise broadly affects the basal ganglia, brain structures involved in motor coordination. Exercise-induced changes in these regions can improve pathological conditions such as Parkinson’s disease and substance use disorders. Importantly, biological sex is a significant factor in the effects of exercise and in the presentation of these basal ganglia-related conditions. Here, we find surprising sex differences in exercise’s influence over motor coordination and neural activity across three extended basal ganglia structures: dorsomedial striatum cholinergic interneurons (CINs), substantia nigra pars compacta (SNc) dopaminergic neurons, and caudal pedunculopontine nucleus (PPN) cholinergic neurons. Using voluntary wheel running, accelerating rotarod, *ex vivo* electrophysiology, and morphological reconstructions, we found that exercise enhances motor coordination, increases SNc excitability, and strengthens excitatory input onto the PPN selectively in female mice. By contrast, exercise increases spontaneous firing rate and reduces dendritic complexity selectively in male CINs. These data reveal sex-specific exercise effects correlated across behavioral and cellular levels.

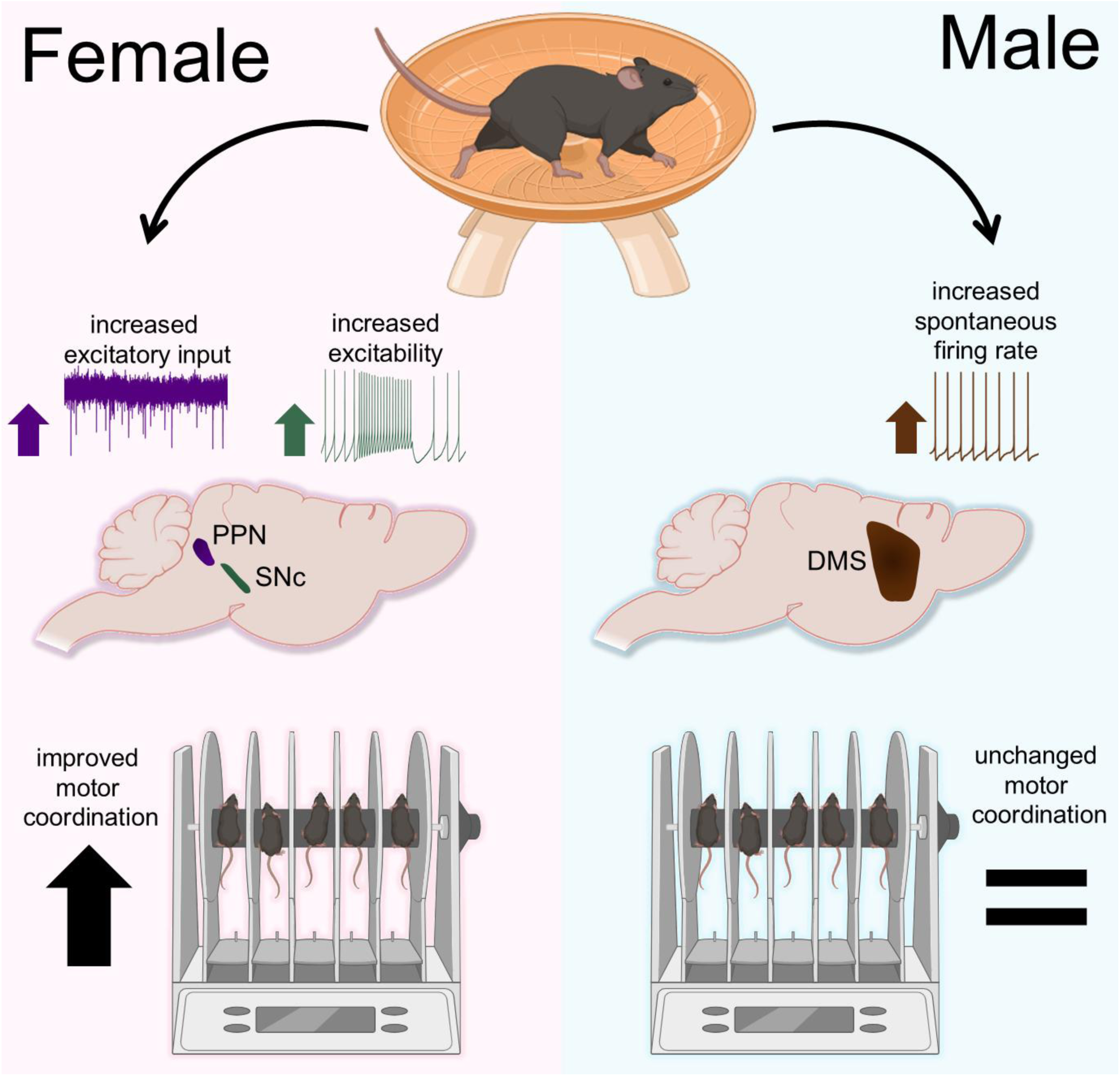

**Highlights:**

- One week of exercise enhances motor coordination in females but not males
- Exercise increases SNc excitability and excitatory input onto the PPN in females
- Exercise increases dorsomedial striatal cholinergic neurons activity in males
- Baseline sex differences in morphology of striatal cholinergic and SNc neurons

## Introduction

Aerobic exercise has many beneficial effects on the brain, including improving motor learning and coordination^1–6^, boosting mood and reducing depression^7–9^, improving sleep quality^10–12^, combating substance use disorders^13–15^, and reducing risk and symptoms of neurodegenerative disorders such as Parkinson’s disease (PD)^16–22^. Each of the above-listed processes involves changes in the extended basal ganglia, which govern motor function, reward processing, and arousal.

There are well-established sex differences both in the susceptibility to neurological disorders and benefits of exercise. Some affective disorders, including depression, are more common in women than men ^23^. On the other hand, PD has a greater incidence in men than women^24,25^. PD symptoms also present differently in men and women, with men generally exhibiting more severe motor deficits such as rigidity, and women exhibiting more extensive nonmotor symptoms^26–28^. Men and women exhibit differential metabolic changes with exercise^29,30^, and women obtain greater cognitive, mood, and cardiovascular benefits from exercise than men^31–35^. Furthermore, the efficacy of exercise in improving basal ganglia-related conditions differs between men and women. For example, women more readily experience benefits of exercise in the context of sleep quality and substance use disorders^10,11,13,15,36^.However, a comprehensive evaluation of the interactions between sex and exercise on physiological properties of multiple modulatory neural types across the extended basal ganglia has not been conducted.

Three major modulatory neuronal populations in the extended basal ganglia, the substantia nigra *pars compacta* (SNc) dopaminergic neurons, pedunculopontine nucleus (PPN) cholinergic neurons, and dorsomedial striatal cholinergic interneurons (CINs) are involved in motor coordination and are affected in PD^37–47^. Pathological activity in these structures contribute to PD motor symptoms^40,48–50^. Exercise alters gene expression^51–55^, inflammatory markers^56–60^, and dopamine dynamics^61–65^ within the these structures. Importantly, while PPN cholinergic neurons have been identified as a critical link between aerobic exercise and enhanced motor coordination^55^, the effects of exercise on their physiological characteristics has not been tested.

Here, we evaluate the broad-ranging effects of exercise across the extended basal ganglia at the cellular level. We find surprising sex differences in exercise’s influence over both motor coordination and neural activity. Using voluntary wheel running, accelerating rotarod, *ex-vivo* whole-cell electrophysiology, and 3D dendritic morphological reconstructions, we found that exercise enhances motor coordination, increases SNc excitability, and strengthens excitatory input onto PPN (but not striatal) cholinergic neurons selectively in female mice. Interestingly, we found that in females, excitatory input to PPN cholinergic neurons was strongest in the mice that ran the most. By contrast, exercise increases spontaneous firing rate and reduces dendritic complexity of dorsomedial striatal CINs selectively in male mice. These sexually-dimorphic behavioral and plasticity changes show that, while exercise has global effects across the brain, it can trigger highly precise circuit alterations in dopaminergic and cholinergic systems of the extended basal ganglia.

## Results

### One week of voluntary running improves motor coordination selectively in female mice

Voluntary wheel running can improve motor coordination in rodents^55,59,62,66,67^, but the interactions between exercise and biological sex have not been elucidated. To test the effect of one week of voluntary wheel running on motor coordination in male and female mice, we provided a saucer-style running wheel in each mouse’s home cage for one week and evaluated subsequent motor performance on the accelerating rotarod (Fig. 1A). Sedentary control mice were given a locked wheel. Mice in the exercise group reliably adopted home-cage wheel running (Fig. 1B). Wheels were removed from the home cages on the first day of rotarod testing, designated day 1. Surprisingly, we found that one week of exercise improved motor coordination only in female mice. The female exercise group exhibited longer latencies to fall off the rotarod than the sedentary group on day 1 (day 1 average latency to fall *N*=17 Sedentary: 143.53 ± 3.96 s, *N*=15 Exercise: 169.74 ± 6.19 s, *t*=-3.57, *p*=0.0015; Fig. 1D, S1D). By contrast, male mice performed at similar levels regardless of whether they exercised or were sedentary (day 1 average latency to fall *N*=13 Sedentary: 150.17 ± 6.13 s, *N*=15 Exercise: 144.67 ± 5.95 s, *t*=0.32, *p*=0.53; Fig. 1E, S1D). Importantly, we found that running behavior did not differ between male and female mice (total distance run, *N*=21 Female: 47.22 ± 4.62 km, *N*=15 Male: 45.09 ± 4.39 km, *t*=0.33, *p*=0.74; distance run on final night, Female: 8.23 ± 0.77 km, Male: 8.11 ± 0.78 km, *t*=0.11, *p*=0.92; Fig. 1C, S1A), indicating that this sex difference in motor enhancement is not explained by females exercising more than males. The motor skill enhancement was also not explained by differences in mouse weight, as exercising and sedentary female mice did not differ in weight at the time of rotarod testing (weight on day 1 *N*=7 Sedentary: 19.86 ± 0.40 g, *N*=7 Exercise: 19.43 ± 0.69 g, *U*=0.91, *p*=0.91; Fig. S1C). However, the male exercise group did weigh less than the male sedentary group (weight on day 1 *N*=7 Sedentary: 25.57 ± 0.90 g, *N*=7 Exercise: 23.14 ± 0.63 g, *t*=2.21, *p*=0.049; Fig. S1C). Improved motor coordination after exercise in females was not long-lasting. The female exercise group showed a trend toward enhanced performance on this task on day 2 after running wheels were removed from the home cages (day 2 average latency to fall, *N*=17 Sedentary: 149.39 ± 6.91 s, *N*=15 Exercise: 169.09 ± 7.66 s, *t*=-1.91, *p*=0.066; Fig. S1D). However, no difference was found between exercise and sedentary female mice the next week (day 9 average latency to fall, Sedentary: 159.61 ± 12.80 s, Exercise: 165.10 ± 7.72 s, *t*=-0.37, *p*=0.71). This indicates that one week of voluntary running in females produces an acute improvement in motor coordination, rather than improving long-term motor learning or recall. Together, these data show that one week of exercise transiently improves motor coordination in female, but not male mice.

**Figure 1:**
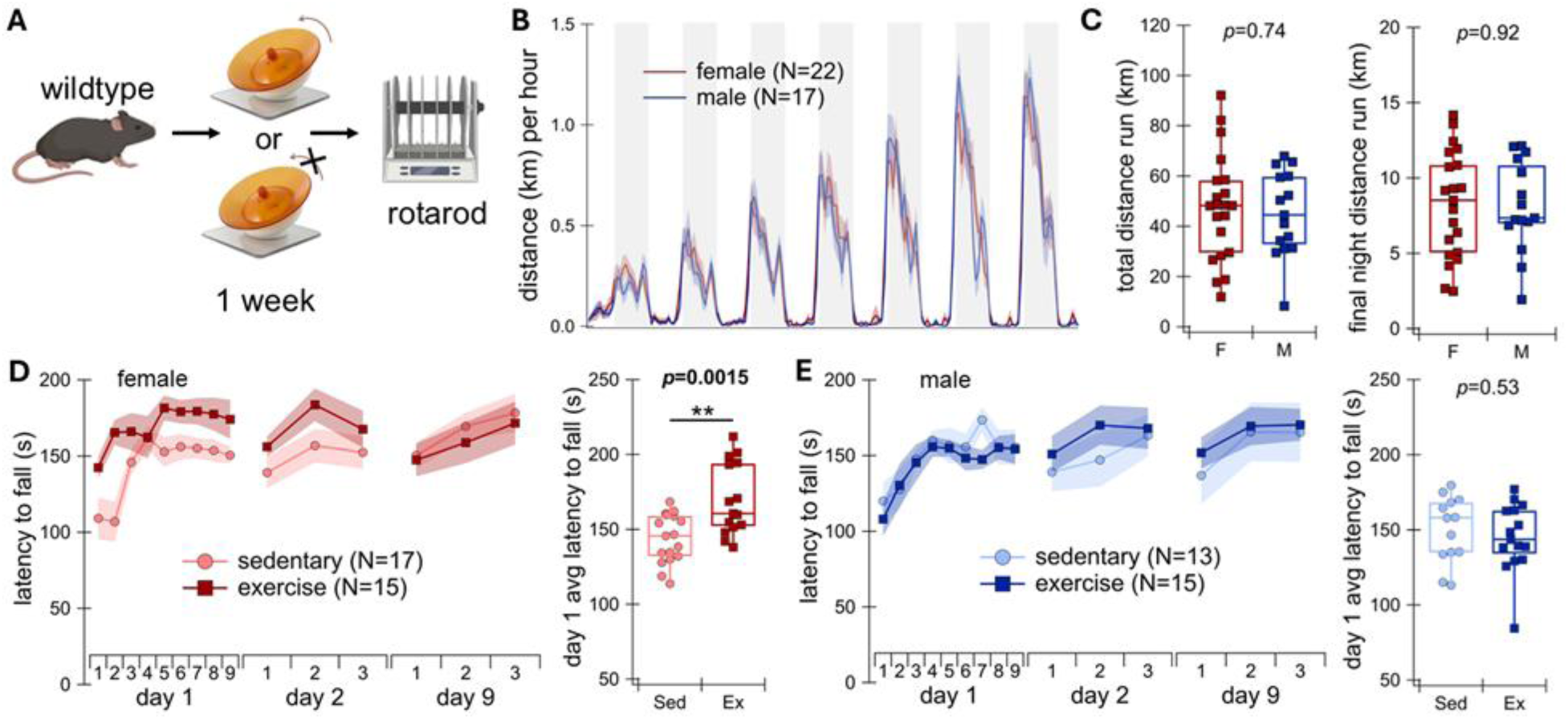
One week of voluntary running improves motor coordination selectively in female mice. **(A)** Schematic of experimental design showing mice given a moving or locked wheel for one week and then tested on the accelerating rotarod task. **(B)** Distance mice ran per hour separate by sex. Shaded regions indicate dark hours, 6pm–6am. **(C)** Total distance run (left) and distance run on the final night (right) compared between females (F) and males (M). **(D)** Left, latency to fall from the rotarod during each trial across the three days of rotarod testing in females. Right, average latency to fall across the 9 trials of day 1 of rotarod testing, compared across sedentary (Sed) and exercise (Ex) female mice. **(E)** Same as in (D) but in male mice. In graphs, data are represented as mean ± SEM. \*\**p* < 0.01. See also Figure S1.

### Voluntary running increases SNc dopaminergic neuron excitability selectively in females

The substantia nigra *pars compacta* (SNc) dopaminergic neurons are important for modulating motor output signals and motor learning^68–70^. Exercise increases dopamine release in the dorsal striatum and protects SNc dopaminergic neurons from degeneration^52,61,62,64,65,71–76^. However, the effects of exercise on SNc physiology and somatodendritic morphology are not known.

To determine whether exercise alters SNc intrinsic properties, we performed whole-cell patch clamp recordings in SNc dopaminergic neurons in DAT-cre/Ai9tdTomato mice after one week of voluntary running (Fig. 2A). Brains were extracted for electrophysiology in the morning after the seventh night of running, which corresponds to the day on which motor coordination benefits were observed in females (Fig. 1D). To test exercise effects on intrinsic excitability, we injected a series of ascending depolarizing currents into SNc neurons. A subset of SNc dopaminergic neurons entered depolarization block, in which sodium channels do not recover from inactivation^77^. Exercise increased the propensity for SNc neurons to enter depolarization block in females (probability of depolarization block at 100 pA injection *n*=50 cells from 7 Sedentary mice: 10.00%, *n*=55 cells from 7 Exercise mice: 27.27%, Fisher’s exact test, *p*=0.0032; Fig. 2E). Exercise female mice showed higher maximum AP frequencies during current injection than sedentary mice (maximum AP frequency at 100 pA injection *n*=51 cells from 7 Sedentary mice: 15.98 ± 1.13 Hz, *n*=54 cells from 7 Exercise mice: 21.09 ± 1.86 Hz, *U*=1778, *p*=0.0098; Fig. 2F, 2J). Exercise did not change input resistance in female mice (input resistance *n*=50 cells from 7 Sedentary mice: 317.26 ± 23.55 MΩ, *n*=58 cells from 7 Exercise mice: 287.24 ± 16.29 MΩ, *U*=1553, *p*=0.53; Fig. 2K), showing that changes in induced AP frequencies are not attributable to changes in membrane depolarization at each current step, but rather that higher frequencies occur at similar membrane voltages. Supporting this idea, we found no significant difference in membrane voltage between groups in the presence of depolarizing or hyperpolarizing currents in SNc neurons of females (Fig. S2A).

**Figure 2:**
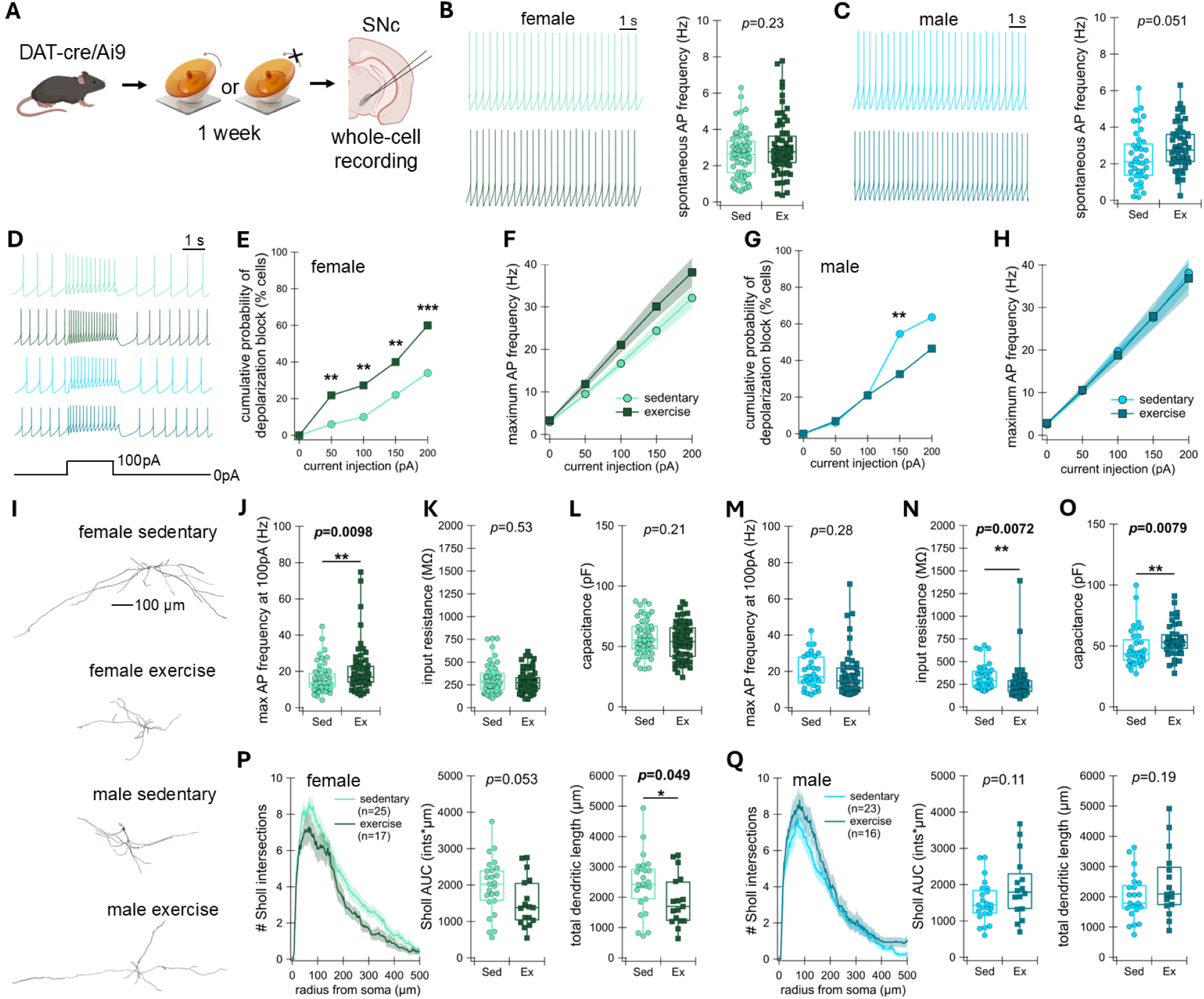
Voluntary running increases intrinsic excitability and decreases dendritic complexity of SNc dopaminergic neurons selectively in females. **(A)** Schematic of experimental design showing mice given a moving or locked wheel for one week and then whole-cell recording in the substantia nigra pars compacta (SNc) dopaminergic neurons in coronal brain slices. **(B)** Left: Example spontaneous membrane voltage traces of SNc neurons in sedentary (top) and exercise (bottom) females. Right: Spontaneous action potential (AP) frequencies in sedentary (Sed) vs exercise (Ex) females. **(C)** Same as in (B) but in male mice. **(D)** Example membrane voltage traces showing before, during, and after 2-s injection of 100 pA. From top to bottom: female sedentary (light green), female exercise (dark green), male sedentary (light blue), male exercise (dark blue). **(E)** Cumulative probability of depolarization block occurring during each current injection step from 0 to 200 pA, increasing in increments of 50 pA, in females. **(F)** Maximum AP frequency in female sedentary and exercise mice during each current step. **(G–H)** Same as in (E–F), but in males. **(I)** Example reconstructed neurons in each of the four groups. **(J)** Maximum AP frequency during the 100 pA current step in females. **(K)** Input resistance in females. **(L)** Capacitance in females. **(M–O)** Same as in J–L, but in males. **(P)** Left: Graph shows number of Sholl intersections at each micron radius from the soma in sedentary and exercise females. Middle: area under the curve (AUC) for each cell’s Sholl graph in females. Right: Total dendritic length in females. **(Q)** Same as in P, but in males. In graphs, data are represented as mean ± SEM. \**p* < 0.05, \*\**p* < 0.01, \*\*\**p* < 0.001. See also Figure S2.

Contrasting with the results in female mice, SNc neurons of males showed decreased excitability after exercise, as evidenced by lower input resistance (*n*=36 cells from 6 Sedentary mice: 331.03 ± 24.06 MΩ, *n*=37 cells from 6 Exercise mice: 276.73 ± 37.43 MΩ, *U*=908, *p*=0.0072; Fig. 2N), and lower changes in voltage in the presence of hyperpolarizing and depolarizing currents (Fig. S2B). Males that exercised also exhibited a decreased likelihood of depolarization block at higher current injections (probability of depolarization block at 150pA injection *n*=33 cells from 6 Sedentary mice: 54.55%, *n*=43 cells from 6 Exercise mice: 33.56%, Fisher’s exact test, *p*=0.0017; Fig. 2G). However, no difference in maximum AP frequency was found between exercised and sedentary males (maximum AP frequency at 100 pA injection *n*=33 cells from 6 Sedentary mice: 19.73 ± 1.58 Hz, *n*=44 cells from 6 Exercise mice: 19.11 ± 1.99 Hz, *U*=620, *p*=0.28; Fig. 2H, 2M). Exercise did not significantly change the spontaneous firing rate of SNc neurons in either females or males, though there was a trend toward increased spontaneous firing rate in exercised male mice (spontaneous AP firing rate *n*=63 cells from 7 Female Sedentary mice: 2.63 ± 0.16 Hz, *n*=63 cells from 7 Female Exercise mice: 3.07 ± 0.20 Hz, *U*=2232, *p*=0.23; *n*=39 cells from 6 Male Sedentary mice: 2.38 ± 0.23 Hz, *n*=49 cells from 6 Male Exercise mice: 2.96 ± 0.18 Hz, *t*=-1.99, *p*=0.051; Fig. 2B–C). Together, these data show that exercise increases SNc intrinsic excitability selectively in female mice.

### Voluntary running decreases SNc dopaminergic dendritic complexity selectively in females

Exercise changes the dendritic morphology of neurons in several brain regions including the dentate gyrus, prefrontal cortex, cerebellum, cuneiform nucleus, dorsal raphe, and ventrolateral medulla^78–82^. To test whether SNc dendritic morphology is altered by exercise, we performed three-dimensional reconstructions of neurobiotin-filled SNc neurons from exercised and sedentary mice and analyzed dendritic complexity. In females, SNc neurons in the exercise group exhibited shorter overall dendritic length (*n*=25 cells from 6 Sedentary mice: 2452.76 ± 193.12 µm, *n*=17 cells from 6 Exercise mice: 1882.32 ± 202.77 µm, *t*=2.037, *p*=0.049; Fig. 2I top, 2P) and trended towards reduced Sholl area under the curve (Sholl AUC *n*=25 cells from 6 Sedentary mice: 1971.12 ± 151.86 Sholl intersections*µm, *n*=17 cells from 6 Exercise mice: 1532.94 ± 164.84 Sholl intersections*µm, *t*=2.00, *p*=0.053; Fig. 2P). Exercise did not change membrane capacitance in SNc neurons of females (*n*=50 cells from 7 Sedentary mice: 57.77 ± 2.12 pF, *n*=58 cells from 7 Exercise mice: 54.04 ± 2.02 pF, *t*=1.27, *p*=0.21; Fig. 2L). Despite these reductions in dendritic complexity, exercise did not decrease the amplitude or frequency of spontaneous excitatory synaptic input onto SNc neurons of females (Fig. S2C). In males, SNc neurons exhibited a higher membrane capacitance in the exercise group (*n*=36 cells from 6 Sedentary mice: 48.40 ± 2.59 pF, *n*=39 cells from 6 Exercise mice: 54.84 ± 2.21 pF, *U*=453, *p*=0.0079; Fig. 2O), indicating larger neurons. However, exercise did not induce dendritic morphology changes in SNc neurons of males (total dendritic length, *n*=23 cells from 5 Sedentary mice: 1996.31 ± 157.30 µm, *n*=16 cells from 4 Exercise mice: 2428.30 ± 277.58 µm, *t*=-1.35, *p*=0.19; Sholl AUC, Sedentary 1523.91 ± 119.26 Sholl intersections*µm, Exercise: 1930.56 ± 215.59 Sholl intersections*µm, *t*=-1.65, *p*=0.11; Fig. 2I bottom, Fig. 2Q). Together these data show that one week of voluntary running decreases SNc neuron dendritic arbors in female, but not male mice.

Overall, these results show sexually dimorphic effects of exercise on SNc dopaminergic neurons: in females, exercise results in less dendritically complex dopaminergic neurons with higher excitability, while in males, exercise results in larger SNc neurons that are less excitable but slightly more spontaneously active.

Baseline sex differences in SNc dendritic complexity have not previously been evaluated. Therefore, we compared the dendritic complexity across sexes in sedentary control mice. We found that females had larger SNc dendritic arbors and higher membrane capacitances than males (Sholl AUC, *n*=25 cells from 6 sedentary Female mice: 1971.12 ± 151.86 # Sholl intersections*µm, *n*=23 cells from 5 sedentary Male mice: 1523.91 ± 119.26 # Sholl intersections*µm, *t*=2.32, *p*=0.025; capacitance, *n*=50 cells from 7 sedentary Female mice: 57.77 ± 2.12 pF, *n*=36 cells from 6 sedentary Male mice: 48.40 ± 2.59 pF, *U*=537, *p*=0.0013; Fig. 3A–B, 3F). While we were not expecting this result, it is not entirely surprising, as estrogen promotes SNc dendritic arborization^83–85^. To our knowledge, these findings represent the first direct comparison of SNc dendritic characteristics in adult male and female mice.

**Figure 3.**
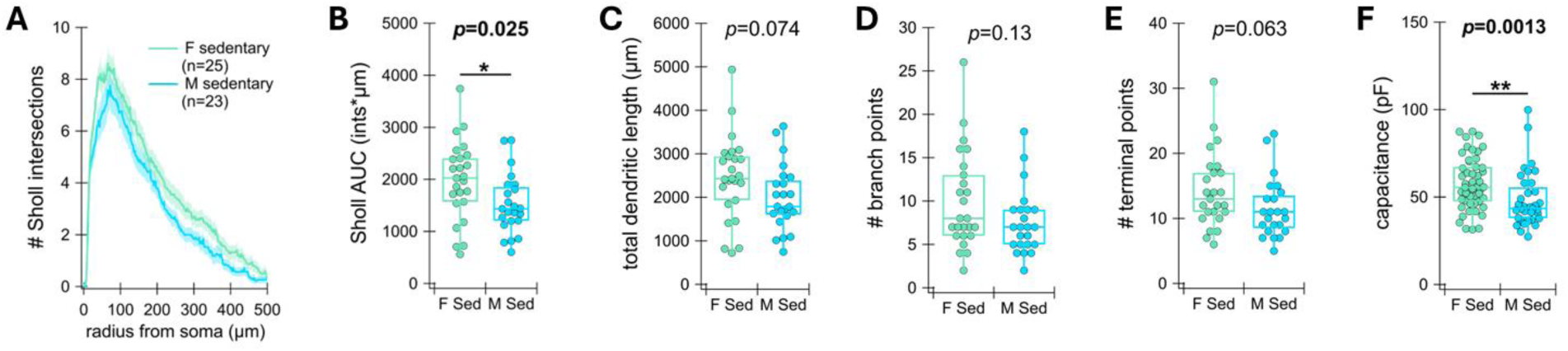
Females have more complex SNc dendritic morphology than males at baseline. **(A)** Number of Sholl intersections at each micron radius from the soma in substantia nigra pars compacta (SNc) dopaminergic neurons of sedentary females (F; green) and males (M; blue) in the same experiment as Figure 2. Data are represented as mean ± SEM. **(B)** Area under the curve (AUC) for each cell’s Sholl graph. **(C)** Total dendritic length (Female: 2452.76 ± 193.12 µm, Male: 1996.31 ± 157.30 µm, *t*=1.83, *p*=0.074). **(D)** Number of dendritic branch points (Female: 9.96 ± 1.12, Male: 7.57 ± 0.78, *U*=214, *p*=0.13). (E) Number of dendritic terminal points (Female: 14.28 ± 1.16, Male: 11.52 ± 0.95, *U*=197, *p*=0.063). **(F)** Cell capacitance. \**p* < 0.05, \*\**p* < 0.01.

### Voluntary running increases dorsomedial striatum cholinergic interneuron spontaneous activity selectively in males

The dorsomedial striatum (DMS) differs from the dorsolateral striatum and mediates early motor learning and supports wheel running behavior^86–94^. Cholinergic interneurons (CINs) in the dorsal striatum play an important part in controlling dopamine release through actions on nicotinic acetylcholine receptors on dopaminergic axons^95–101^. Activity levels and excitability of CINs are altered in PD models^47,102–106^, and modulation of their activity levels contributes to motor skill learning, vigor, and cognitive flexibility^44,107–111^. However, the effect of exercise on these neurons has not yet been determined. Thus, to test the effect of exercise on neuronal properties of CINs in the DMS, we performed whole-cell patch clamp recordings in DMS CINs in ChAT-cre/Ai9-tdTomato mice after one week of voluntary running (Fig. 4A).

**Figure 4:**
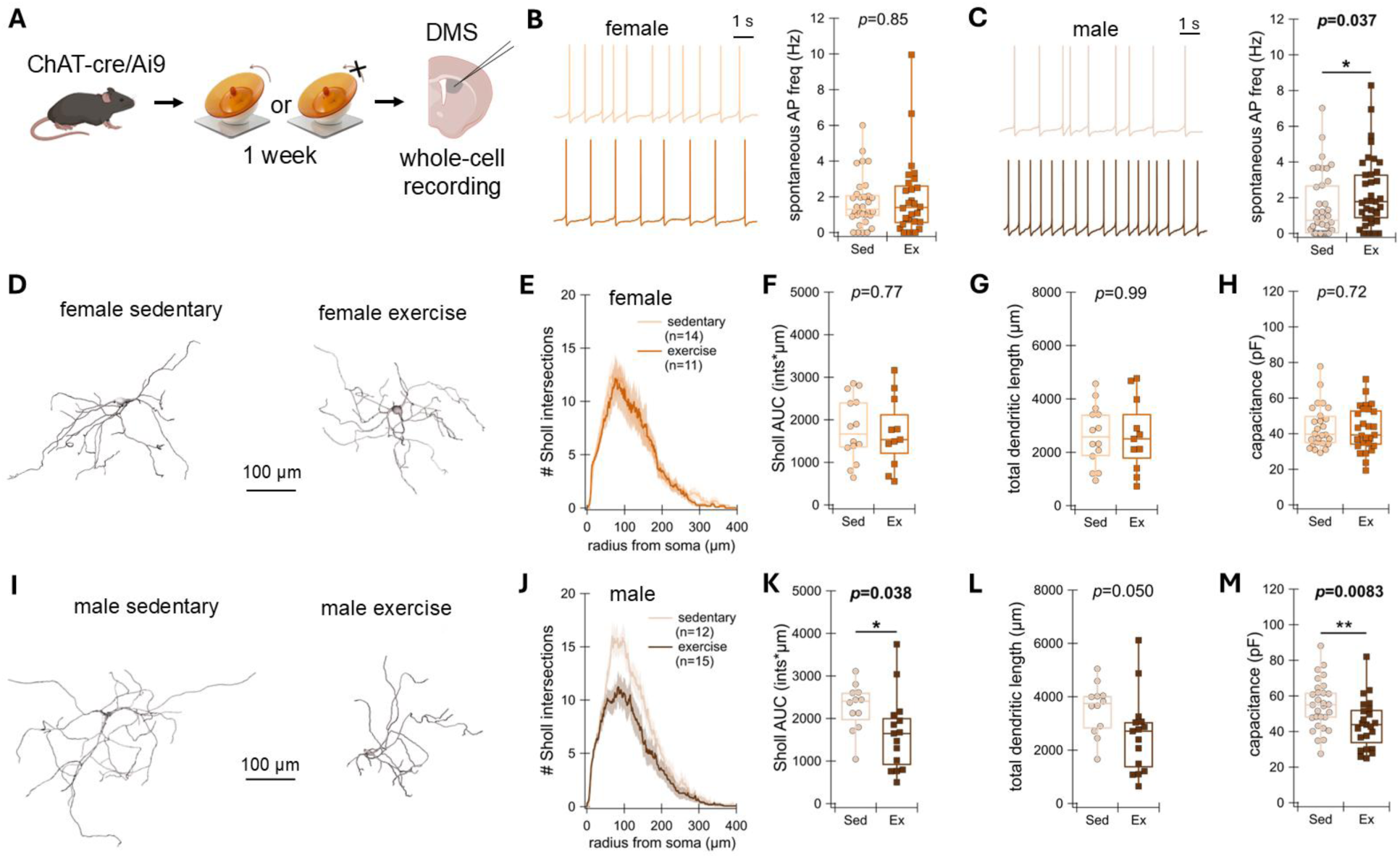
Voluntary running increases spontaneous activity and decreases dendritic complexity of DMS CINs selectively in males. **(A)** Schematic of experimental design showing mice given a moving or locked wheel for one week and then whole-cell recording in dorsomedial striatal (DMS) cholinergic interneurons (CINs) in coronal brain slices. **(B)** Left: Example spontaneous membrane voltage traces of DMS CINs in sedentary (top) and exercise (bottom) females. Right: Spontaneous action potential (AP) frequencies in sedentary (Sed) vs exercise (Ex) females. **(C)** Same as in (B) but in male mice. **(D)** Example reconstructed CINs in sedentary and exercise female groups. **(E)** Graph shows number of Sholl intersections at each micron radius from the soma in sedentary and exercise females. **(F)** Area under the curve (AUC) for each cell’s Sholl graph in females. **(G)** Total dendritic length in females. **(H)** Capacitance in females. **(I–M)** Same as in (D–H) but in males. In graphs, data are represented as mean ± SEM. \**p* < 0.05, \*\**p* < 0.01. See also Figure S3.

Our results show that CINs in male mice that have exercised exhibited higher spontaneous AP firing rates than sedentary males (*n*=38 cells from 8 Sedentary mice: 1.49 ± 0.29 Hz, *n*=37 cells from 7 Exercise mice: 2.24 ± 0.33 Hz, *U*=507, *p*=0.037; Fig. 4C). In females, on the other hand, we observed no difference in spontaneous firing rate (*n*=32 cells from 8 Sedentary mice: 1.91 ± 0.41 Hz, *n*=28 cells from 8 Exercise mice: 1.71 ± 0.26 Hz, *U*=435, *p*=0.85; Fig. 4B). Exercise did not alter CIN excitability (Fig. S3A–B) or excitatory synaptic input (Fig. S3E–F) in either sex. However, exercise caused CINs in females to be more easily hyperpolarized (change in membrane potential at −100 pA injection *n*=28 cells from 8 Sedentary mice: −11.12 ± 0.55 mV, *n*=24 cells from 8 Exercise mice: −13.28 ± 0.80 mV, *U*=227, *p*=0.046; Fig. S3C). Therefore, exercise increases spontaneous activity in DMS CINs in male but not female mice.

### Voluntary running decreases dorsomedial striatum cholinergic interneuron dendritic complexity selectively in males

CIN dendritic arborization is sensitive to dopamine levels, as loss of dopamine neurons in the 6-OHDA model of PD increases CIN dendritic complexity^112,113^. Because exercise increases dorsal striatal dopamine release^61,62,65^, we hypothesized that exercise would decrease CIN dendritic arborization. To test this hypothesis, we reconstructed CINs from mice with and without one week of voluntary running. In male mice, we found that, indeed, CINs in the exercise group exhibited reduced dendritic complexity, as evidenced by lower Sholl AUC (*n*=12 cells from 4 Sedentary mice: 2264.92 ± 159.76 Sholl intersections*µm, *n*=15 cells from 7 Exercise mice: 1650.13 ± 229.68 Sholl intersections*µm, *t*=2.20, *p*=0.038; Fig. 4I–K) and a trend towards smaller total dendritic length (Sedentary: 3546.35 ± 275.16 µm, Exercise: 2579 ± 380.36 µm, *t*=2.06, *p*=0.050; Fig. 4L). The area of reduced complexity occurred particularly within 60–110 µm from the soma. Reduced CIN capacitance in exercised male mice accompanied this change (*n*=29 cells from 6 Sedentary mice: 54.94 ± 2.57 pF, *n*=23 cells from 5 Exercise mice: 44.27 ± 2.89 pF, *t*=2.76, *p*=0.0083; Fig. 4M). No differences in morphological measures were found between the female exercise and sedentary groups (Sholl AUC, *n*=14 cells from 6 Sedentary mice: 1783.50 ± 201.61 Sholl intersections*µm, *n*=11 cells from 5 Exercise mice: 1689.73 ± 250.71 Sholl intersections*µm, *t*=0.29, *p*=0.77; total dendritic length, Exercise: 2628.05 ± 413.42 µm, Sedentary: 2628.77 ± 300.04 µm, *t*=-0.0014, *p*=0.99; capacitance *n*=26 cells from 7 Sedentary mice: 43.75 ± 2.36 pF, *U*=368, *p*=0.77, *n*=27 cells from 8 Exercise mice: 42.53 ± 2.40 pF; Fig. 4D–H).

As in the SNc, we found baseline sex differences in the dendritic arborization of DMS CINs in control mice. CINs of the female sedentary group exhibited lower dendritic lengths and capacitances than those of the male group (total dendritic length, *t*=-2.25, *p*=0.034; capacitance, *U*=566, *p*=0.0012; Fig. 5A, 5C, 5F). Altogether, these findings show that exercise alters electrophysiological and morphological characteristics in DMS CINs selectively in males: exercise makes CINs of males more spontaneously active and reduces their extensive dendritic arborization to a level similar to that found in females.

**Figure 5.**
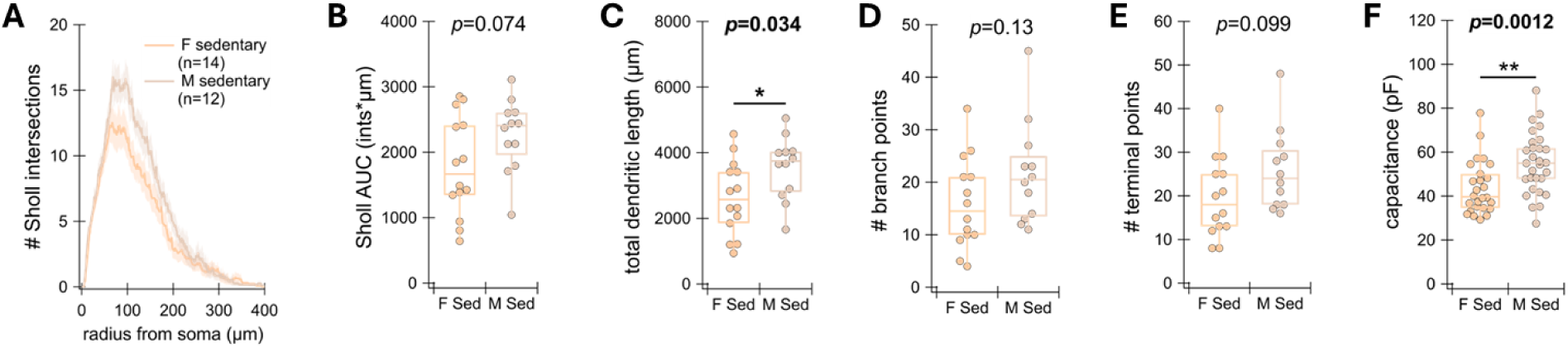
Males have more complex DMS CIN dendritic morphology than females at baseline. **(A)** Number of Sholl intersections at each micron radius from the soma in dorsomedial striatal (DMS) cholinergic interneurons (CINs) of sedentary females (F; orange) and males (M; brown) in the same experiment as Figure 4. Data are represented as mean ± SEM. **(B)** Area under the curve (AUC) for each cell’s Sholl graph (Female: 1783.50 ± 201.61 # intersections*µm, Male: 2264.92 ± 159.76 # intersections*µm, *t*=-1.87, *p*=0.074). **(C)** Total dendritic length. **(D)** Number of dendritic branch points (Female: 15.93 ± 2.32, Male: 21.58 ± 2.81, *t*=-1.55, *p*=0.13). **(E)** Number of dendritic terminal points (Female: 19.57 ± 2.46, Male: 25.83 ± 2.69, *t*=-1.72, *p*=0.099). **(F)** Cell capacitance. \**p* < 0.05, \*\**p* < 0.01.

### Voluntary running reduces PPN cholinergic excitability in females across all estrous stages

The cholinergic neurons of the pedunculopontine nucleus (PPN) are implicated in exercise-driven enhancement of motor learning^55^ as well as the modulation of sleep-wake state^114,115^. Cholinergic neurons of the caudal PPN exhibit an upregulation of *c-Fos* after one week of voluntary running in male mice^55^, suggesting increased PPN neural activity. However, it is not known whether exercise changes PPN intrinsic activity levels, excitability, or synaptic connectivity. Therefore, we tested the effect of exercise on neuronal properties in caudal PPN cholinergic neurons by performing whole-cell patch clamp recording in male and female ChAT-cre/Ai9-tdTomato mice in exercise and sedentary conditions (Fig. 6A). We found that caudal PPN cholinergic neuron spontaneous firing rates did not differ between exercise and sedentary groups in females (*n*=36 cells from 7 Sedentary mice: 4.03 ± 0.61 Hz, *n*=40 cells from 9 Exercise mice: 2.89 ± 0.61 Hz, *U*=891, *p*=0.073; Fig. 6B) or males (*n*=40 cells from 11 Sedentary mice: 4.30 ± 0.69 Hz, *n*=33 cells from 8 Exercise mice: 3.54 ± 0.59 Hz, *U*=823, *p*=0.77; Fig. 6C). However, exercise did decrease caudal cholinergic PPN neuron excitability in females (average AP frequency at 100 pA injection, *n*=28 cells from 7 Sedentary mice: 21.30 ± 2.43 Hz, *n*=32 cells from 9 Exercise mice: 14.80 ± 1.29 Hz, *U*=298, *p*=0.026; input resistance, *n*=29 cells from 6 Sedentary mice: 410.35 ± 36.51 MΩ, *n*=34 cells from 9 Exercise mice: 269.88 ± 17.82 MΩ, *U*=742, *p*=0.00045; Fig. 6D–E), but not males (average AP frequency at 100 pA injection, *n*=42 cells from 11 Sedentary mice: 26.54 ± 3.12 Hz, *n*=29 cells from 8 Exercise mice: 22.88 ± 2.23 Hz, *U*=600, *p*=0.92; input resistance n=39 cells from 9 Sedentary mice: 470.70 ± 51.92 MΩ, n=19 cells from 6 Exercise mice: 538.77 ± 76.13 MΩ, *U*=307, *p*=0.30; Fig. 6D, 6F).

**Figure 6:**
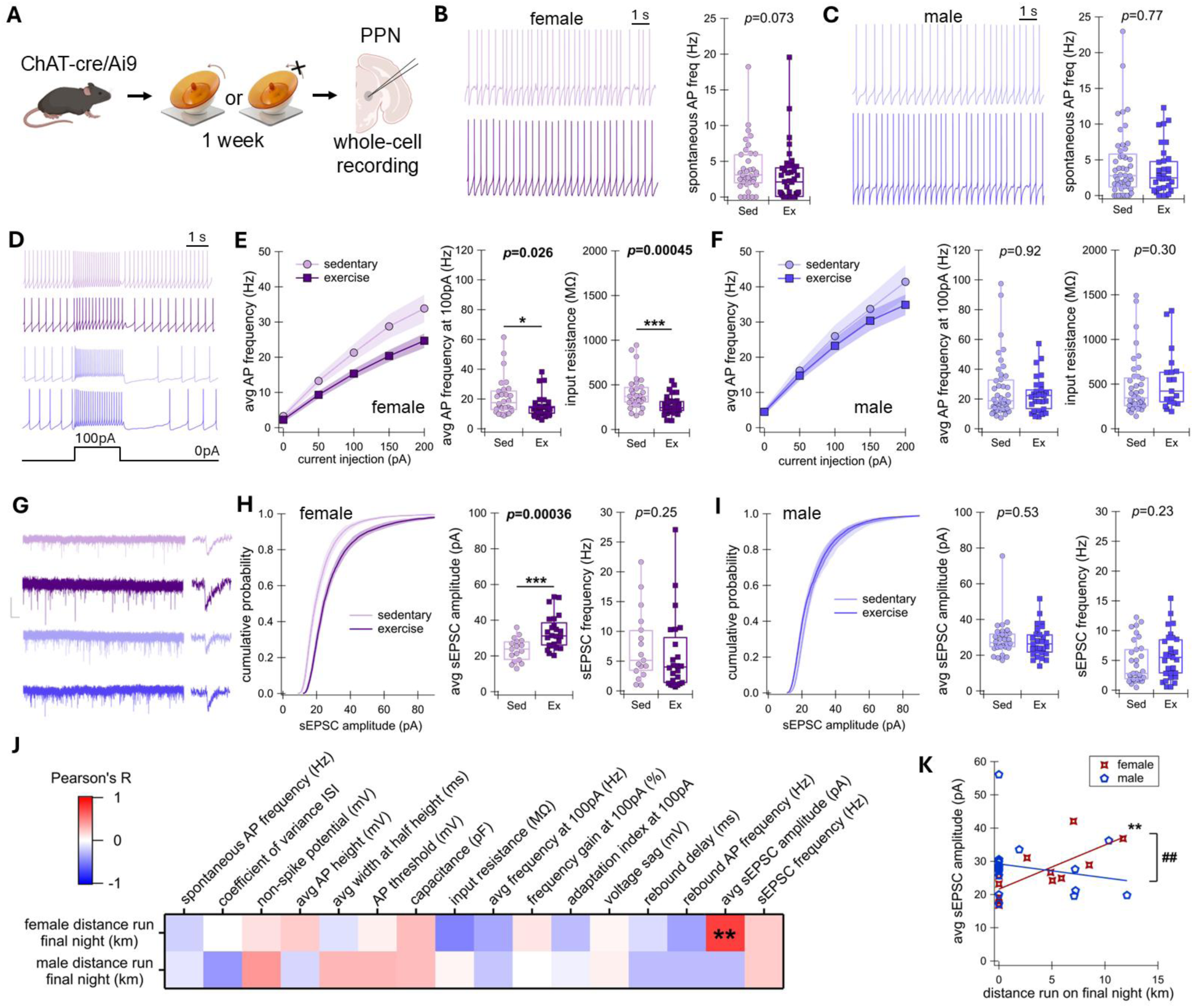
Voluntary running reduces PPN cholinergic neuron intrinsic excitability and increases excitatory synaptic input selectively in female mice. **(A)** Schematic of experimental design showing mice given a moving or locked wheel for one week and then whole-cell recording in caudal pedunculopontine nucleus (PPN) cholinergic neurons in coronal brain slices. **(B)** Left: Example spontaneous membrane voltage traces of PPN cholinergic neurons in sedentary (top) and exercise (bottom) females. Right: Spontaneous action potential (AP) frequencies in sedentary (Sed) vs exercise (Ex) females. **(C)** Same as in (B) but in male mice. **(D)** Example membrane voltage traces showing before, during, and after 2-s injection of 100 pA. From top to bottom: female sedentary (light pink), female exercise (dark pink), male sedentary (light purple), male exercise (dark purple). **(E)** Left: Average AP frequency in female sedentary and exercise mice during each current injection step from 0 to 200 pA, increasing in increments of 50pA. Middle: Average AP frequency during the 100 pA current step in females. Right: Input resistance in females. **(F)** Same as in (E) but in males. **(G)** Left: Example traces showing current during 10-s holding at −70mV for all four groups. Scale bars: 0.5 sec horizontal, 50 pA vertical. Right: Example spontaneous excitatory postsynaptic currents (sEPSCs). **(H)** Left: Cumulative probability of sEPSC amplitude in females. Middle: Average sEPSC amplitude in females. Right: sEPSC frequency in females. **(I)** Same as in (H), but in males. **(J)** Heat map showing correlations between distance run on the final night and the averaged electrophysiological property values in each mouse. Warmer colors indicate positive correlations and colder colors indicate negative correlations. **(K)** Each mouse’s averaged sEPSC amplitude plotted against the mouse’s distance run on the final night among males and females. ## indicates significant difference between relationships in males vs females. In graphs, data are represented as mean ± SEM. \**p* < 0.05, \*\**p* < 0.01, \*\*\**p* < 0.001, ##*p* < 0.01. See also Figures S4 and S5.

Because hormone fluctuations across estrous stages can influence neural activity^116–119^ and may interact with exercise effects^120^, we used vaginal cytology to track estrous stage in female mice after euthanasia for electrophysiology. The exercise-dependent decrease in current-frequency curve occurred in estrus, metestrus, and diestrus (Fig. S4). Insufficient sample size prevented this comparison for proestrus. Overall, these data show that exercise reduces caudal cholinergic PPN excitability in female mice across the estrous cycle.

### Amount of running correlates with strengthened excitatory synaptic input onto PPN cholinergic neurons selectively in females

Finding a decrease in PPN cholinergic neuron excitability after exercise is surprising because lower excitability would not be expected to correspond to an increase in immediate early gene expression (e.g., *c-Fos*) as previously observed^55^. However, another mechanism by which increased neuronal activity (and therefore *c-Fos* expression) could be driven is through the strengthening of excitatory synaptic input onto these neurons. Measuring spontaneous excitatory post-synaptic currents (sEPSCs), we found that one week of voluntary running increased the strength of excitatory synaptic input to caudal PPN cholinergic neurons selectively in female mice. The female exercise group showed larger average sEPSC amplitudes than sedentary (*n*=17 cells from 6 Sedentary mice: 23.46 ± 1.58 pA, *n*=24 cells from 8 Exercise mice: 33.25 ± 1.94 pA, *t*=-3.90, *p*=0.00036; Fig. 6G–H). Importantly, excitatory current amplitude correlated with amount of exercise in females (distance run on final night vs mouse average sEPSC amplitude, *N*=13 mice, Pearson’s *r*=0.73, *p*=0.0049; Fig. 6J–K, Fig. S5M).

Excitatory input did not differ between male exercise and sedentary groups (average sEPSC amplitude, *n*=32 cells from 12 Sedentary mice: 29.10 ± 1.79 pA, *n*=29 cells from 11 Exercise mice: 27.43 ± 1.57 pA, *U*=519, *p*=0.43; Fig. 6G, 6I). We also did not find a significant correlation between EPSC amplitude and amount of exercise in male mice (distance run on final night vs mouse average sEPSC amplitude, *N*=18 mice, Pearson’s *r*=-0.20, *p*=0.43; Fig. 6J–K; Fig. S5N). The relationship between distance run and sEPSC amplitude differed between females and males (*z*=2.76, *p*=0.0029; Fig. 6K).

Curiously, PPN cholinergic neurons of males and females also differed in excitatory input at baseline, where sedentary males exhibited larger sEPSC amplitudes than sedentary females (average sEPSC amplitude, *n*=17 cells from 6 Female mice: 23.46 ± 1.58, *n*=32 cells from 12 Male mice: 29.10 ± 1.79, *U*=166, *p*=0.026; Fig. 6G, Fig. 7). These data show that exercise strengthens excitatory synaptic input onto caudal cholinergic PPN neurons in female, but not male mice.

**Figure 7.**
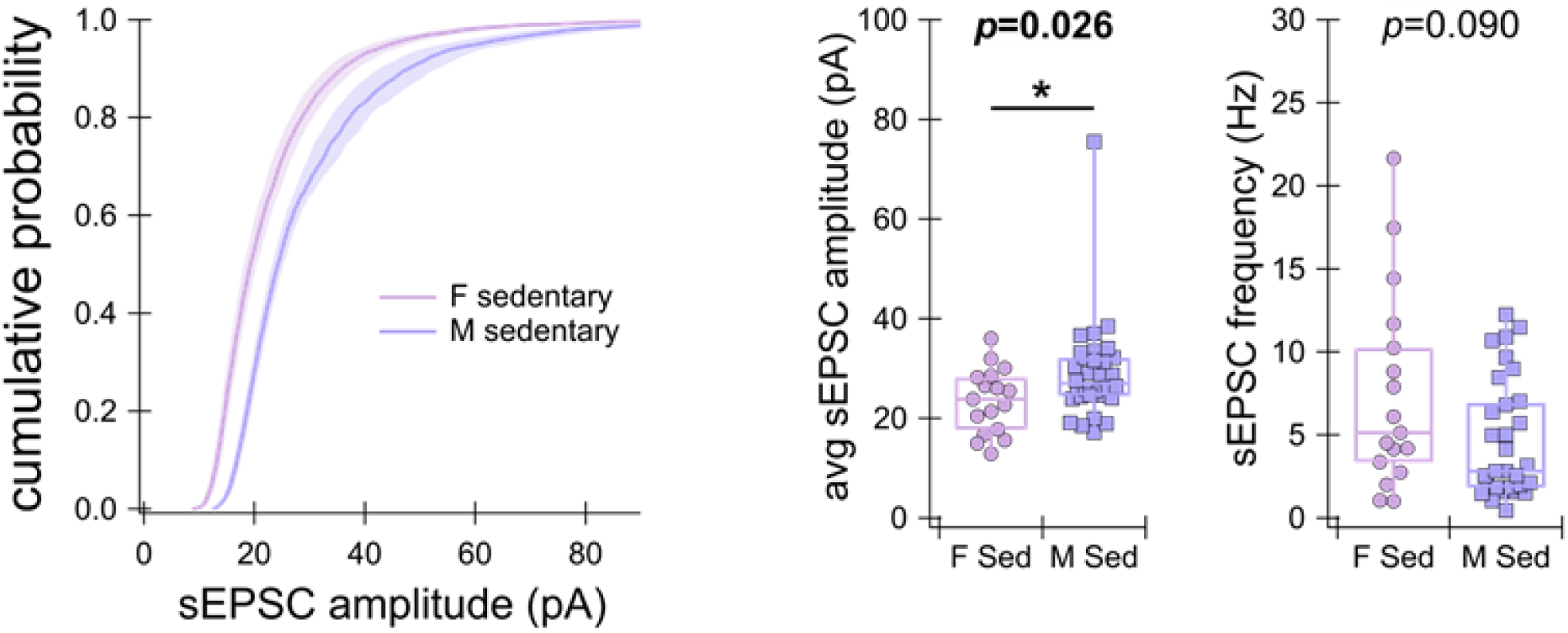
PPN cholinergic neurons in males receive higher amplitudes of spontaneous excitatory input than in females at baseline. Left: Cumulative probability of spontaneous excitatory postsynaptic current (sEPSC) amplitude in caudal pedunculopontine nucleus (PPN) cholinergic neurons of sedentary females (F; pink) and males (M; purple) in the same experiment as Figure 6. Data are represented as mean ± SEM. Middle: Average sEPSC amplitude. Right: sEPSC frequency (Female: 7.43 ± 1.44 Hz, Male: 4.59 ± 0.63 Hz; *U*=353, *p*=0.090). \**p* < 0.05.

No morphological differences due to exercise were found in PPN cholinergic neurons of either sex (Sholl AUC in females, *n*=8 cells from 5 Sedentary mice: 1162.37 ± 231.13 Sholl intersections*µm, *n*=14 cells from 5 Exercise mice: 1050.50 ± 168.01 Sholl intersections*µm, *U*=60, *p*=0.82; Sholl AUC in males, *n*=17 cells from 6 Sedentary mice: 1272.65 ± 154.80 Sholl intersections*µm, *n*=19 cells from 7 Exercise mice: 1375.21 ± 107.24 Sholl intersections*µm, *U*=132, *p*=0.36; Fig. S5C–L). There were also no baseline differences in PPN cholinergic neuron morphology between male and female sedentary mice (Sholl AUC, *U*=71, *p*=0.89). Altogether, these data show that exercise increases excitatory input but decreases excitability in caudal PPN cholinergic neurons selectively in females.

## Discussion

Here we show that one week of voluntary running has sex-specific effects on motor coordination and on the intrinsic properties of SNc dopaminergic neurons, DMS CINs, and caudal PPN cholinergic neurons. Overall, these findings give important insight into how the global effects of exercise result in precise changes to extended basal ganglia physiology and how these effects depend on biological sex.

### Baseline sex differences in the extended basal ganglia

We found baseline sex differences in dendritic morphology of SNc dopaminergic neurons and DMS CINs. Many physiological differences have been found between males and females in the dorsal striatum^121^. For instance, at baseline, female rodents have higher stimulated dopamine release amplitudes as well as higher rates of dopamine clearance than males^122–127^. Despite these extensive dopamine axon-localized differences, spontaneous activity and other electrophysiological properties of the SNc neurons themselves do not differ between male and female mice until over one year of age^128^. Our data in adult mice are consistent with no sex-differences in spontaneous SNc activity. However, in support of baseline sex differences in the SNc, we have found larger capacitances and dendritic arborization in female vs male mice (Fig. S3). Finding sex differences in dendritic arborization is especially important because it raises the possibility that the integration of afferent signals may differ in these neurons between sexes, potentially influencing SNc neuron output^129^.

Previous work on sex differences in dorsal striatal CINs has found similar intrinsic properties across sexes and across female estrous stages^130^. However, female mice have fewer ChAT-expressing neurons in the dorsal striatum than males, though, curiously, females have more ChAT *mRNA*-expressing cells^131,132^. Despite minimal physiological differences in CINs, behavioral differences between male and female mice have been observed after various CIN manipulations^133–135^. Similar to Kövesdi et al., we found no sex differences in neurophysiological properties. However, we find that CINs of males have greater dendritic complexity and membrane capacitance than CINs of females. CIN dendritic arborization is sensitive to dopamine depletion models of PD and genetic models of Huntington’s disease^112,113,136^. Thus, our results raise the possibility that sex differences may play a role in the CIN response in these models.

Sex differences are under-explored in the PPN. One study has shown that PPN cholinergic neurons increase in activity in response to ethanol treatment in male but not female mice^137^. We find that PPN cholinergic neuron excitability and strength of excitatory input are more sensitive to exercise in females compared to males. Additionally, this increased sensitivity does not change across the estrous cycle (Fig. S4), indicating that exercise effects on intrinsic properties of these neurons are not dependent on acute ovarian hormone levels on the day of recording, but rather are dependent on more long-term differences between sexes such as general hormonal and/or gene expression patterns. Our finding that males exhibit greater sEPSC amplitudes than females at baseline (Fig. 7) calls for greater attention to be given to sex differences in excitatory afferents to the PPN. The current evidence for notable differences in PPN cholinergic neuron response to stimuli (exercise and ethanol) highlights the need for more investigation into functional implications for sex differences in the PPN.

### Sex differences in neural effects of exercise

Our results indicate that there are sex-specific effects of exercise within the extended basal ganglia, and these effects may explain sex differences in exercise effects on related behaviors and disorders. As impairment of dopaminergic systems contribute to depression and other mood disorders^138–140^, the exercise-driven increase in SNc neuron excitability that we observe in females may contribute to the exercise improvement of motivation and reduction of anhedonia^9^. The sex specificity of our results may help explain findings from preclinical studies showing that females more readily show resistance to stress-induced anhedonia after exercise^141–144^.

Evidence suggests that women obtain greater benefits of exercise than men on sleep quality, including its effects on rapid eye movement and slow wave sleep phases^10,11,36^. Despite the established role of PPN cholinergic neurons in arousal, sleep-wake state, and rapid eye movement sleep through involvement in the ascending reticular activating system^114,115,145^, research is lacking on the potential for these neurons to be involved in the connection between exercise and improved sleep quality. The improved signal to noise ratio that we observe in PPN cholinergic neurons may allow for more efficient cholinergic communication to downstream brain structures such as the thalamus, but future research is needed to determine whether this could contribute to sex differences in exercise-mediated sleep improvements.

### Implications for motor coordination

We find that female mice exhibit enhanced motor coordination after one week of voluntary running, while male mice do not. Women have greater sensitivity to other physiological effects of exercise such as cardiovascular benefits^32,33^. Exercise-induced increases in the levels of brain-derived neurotrophic factor may also be stronger in females, as determined through meta-analysis of preclinical studies^146^. Nevertheless, aerobic exercise can improve motor function in healthy males^2,3,55,62,66^. One explanation for the sex differences we observe after one week of exercise could be that male mice require more exercise to receive the same benefits as female mice.

Our findings contrast with a previous study investigating exercise and rotarod performance^55^. Specifically, this study found an improvement in motor skill *learning* in male mice, with both exercise and sedentary groups starting at comparable rotarod *performance*. They did not test female mice. In our study, while male mice do start at similar rotarod performances, the exercise group does not show any enhancement in motor learning. Our female mice, on the other hand, show an immediate enhancement of motor coordination after exercise. Comparing methodology between our study and the Li and Spitzer study, we use the same mouse genetic background, wheel style, rotarod settings, and session parameters (number, frequency, duration). It is possible that protocol differences such as circadian period during which rotarod was performed (light hours vs dark hours), control condition (locked wheel vs no wheel), or unaccounted for environmental conditions altered behavioral outcomes between studies.

Along with female-specific enhancement of motor coordination, we also found female-specific exercise effects in two key neural populations that are involved in motor function: SNc dopaminergic neurons and PPN cholinergic neurons. This sex specificity points to these neural changes as potential contributors to improvements in motor coordination. One neural change of note is the increase in excitability of SNc neurons in female mice. Higher maximal firing rates may facilitate greater phasic dopamine release, which is important for motor and reward learning^69,147–149^. We observed changes in DMS cholinergic interneuron spontaneous firing and dendritic complexity due to exercise, but only in male mice. As these mice belong to the population that did not exhibit enhancement of motor coordination, we predict that these changes do not underlie this motor enhancement, and that improvement in motor coordination in females does not involve changes in intrinsic properties of DMS CINs. Another female-specific neural change that may drive enhanced motor coordination is increased excitatory input strength onto caudal PPN cholinergic neurons in combination with a decrease in intrinsic excitability. Together, these changes suggest homeostatic plasticity, in which larger excitatory input would result in a compensatory downregulation of intrinsic excitability (or vice versa). The result of these changes could make caudal cholinergic PPN neurons more discriminatory towards the potentiated excitatory inputs by increasing signal to noise ratio. These inputs could come from a number of regions that send excitatory projections to PPN cholinergic neurons, including the cortex, superior colliculus, hypothalamus, and subthalamic nucleus^150^. More work is needed to determine which inputs to caudal PPN cholinergic neurons are potentiated by exercise.

### Implications for Parkinson’s disease

In addition to the well-established ability for exercise to improve symptoms in PD patients^16,17,20,22,151^, exercise also can reduce the risk of developing PD in both men and women^19,21,152^. These protective effects have been demonstrated through toxin PD rodent models, which show that pretreatment with an exercise regimen can reduce the loss of SNc dopaminergic neurons^53,153–156^. Mechanisms underlying the protective effects of exercise on SNc neurons include improved mitochondrial function and reduced oxidative stress. Our finding that exercise increases SNc neuron excitability without increasing spontaneous firing rates in females suggests that exercise may optimize dopaminergic neuron firing during motor tasks without increasing basal calcium influx. This scenario could support motor coordination without increasing tonic mitochondrial load. Additionally, the exercise-induced reduction in dendritic complexity that we observed in females may be protective by reducing cellular metabolic requirements. Future work is needed to determine whether the effects of exercise on the dopaminergic system could contribute to the reduced prevalence of PD in women compared to men.

## STAR Methods

### KEY RESOURCES TABLE

| REAGENT or RESOURCE | SOURCE | IDENTIFIER |
| --- | --- | --- |
| Antibodies |  |  |
| Streptavidin Cy5 | Invitrogen | Cat#: SA1011 |
| Deposited data |  |  |
| Raw and analyzed data | This paper |  |
| Experimental models: Organisms/strains |  |  |
| C57BL/6J | Jackson | RRID: IMSR_JAX:000664 |
| ChAT-cre mice: B6.129S-Chat<sup>tm1(cre)Lowl/MwarJ</sup> | Jackson | RRID: IMSR_JAX:031661 |
| DAT-cre mice: B6.SJL-Slc6a3<sup>tm1.1(cre)Bkmn/J</sup> | Jackson | RRID: IMSR_JAX:006660 |
| Ai9-tdTomato mice: B6.Cg-Gt(ROSA)26Sor<sup>tm9(CAG-tdTomato)Hze/J</sup> | Jackson | RRID: IMSR_JAX:007909 |
| Software and algorithms |  |  |
| Igor Pro | WaveMetrics | Version 9.0.5.1 |
| Wheel Manager | Med Associates, Inc | Version 2.04 |
| Clampex | Molecular Devices | Version 11.2 |
| Leica Application Suite X | Leica Microsystems | Version 3.5.6 |
| Imaris | Oxford University | Version 10.1.0 |
| BioRender | Science Suite, Inc. |  |
| Other |  |  |
| Wireless running wheel for mice | Med Associates, Inc | ENV-047 |
| Rotarod for mice | Ugo Basile | 47650 |
| Monocular LED student compound microscope | AmScope | D130 Series |
| Upright differential interference contrast microscope | Evident | OpenStand |
| Vibratome | Leica | VT1000S |
| Pipette puller | Sutter Instrument | P-97 |
| Amplifier | Multiclamp | 700B |
| Digitizer | Axon | Digidata 1550B |
| Laser confocal microscope | Leica | SP8 |

### RESOURCE AVAILABILITY

#### Lead contact

Further information and requests for resources and reagents should be directed to and will be fulfilled by the lead contact, Dr. Rebekah C. Evans.

#### Materials availability

This study did not generate new unique reagents.

#### Data and code availability

All data and code supporting the current study will be shared by the lead contact upon request.

The morphological data collected in this study is available on NeuroMorpho.org and is publicly available as of the date of publication.

Any additional information required to reanalyze the data reported in this paper is available from the lead contact upon request.

### EXPERIMENTAL MODEL AND STUDY PARTICIPANT DETAILS

#### Animal welfare

All animal handling, procedures, and care were performed in accordance with regulations set by the Institutional Animal Care and Use Committee (IACUC) at Georgetown University Medical Center. Measures were taken to prevent animal discomfort and to minimize animal numbers.

#### Animal subjects

Adult male and female mice of both sexes with a C57Bl/6 background were used for all experiments. Mice were housed in a standard 12-hour light/12-hour dark cycle and had *ad libitum* access to food and water. Wildtype mice (p55–p131) were used for accelerating rotarod experiments. Adult ChAT-cre/Ai9-tdTomato and DAT-cre/Ai9-tdTomato mice (p46–p247) were created by breeding ChAT-cre homozygous with Ai9 homozygous mice and were used for electrophysiological experiments. Age-matched littermates were assigned randomly to either the exercise or the sedentary condition. Rotarod experiments were conducted during the light cycle.

#### Experimental groups

For running wheel accelerating rotarod experiments, 32 female and 28 male wildtype mice were used. Electrophysiology experiments in cholinergic neurons used 19 female and 26 male ChAT-cre/Ai9-tdTomato mice. The experiments in dopaminergic neurons used 14 female and 12 male DAT-cre/Ai9-tdTomato mice.

### METHOD DETAILS

#### Voluntary running

For the accelerating rotarod experiments and electrophysiology experiments, mice were single-housed with a Med Associates wireless saucer-style running wheel for 7 days. Sedentary control mice were given a locked wheel preventing rotation. Wheel tracking data in units of kilometers run per hour was recorded for a subset of mice using Wheel Manager software. One mouse in the exercise group was excluded from analysis due to not running for the first four nights. Running amounts did not differ between the wildtype mice used in the accelerating rotarod task and the two genotypes used for whole-cell patch clamp experiments (Fig. S1B).

#### Accelerating rotarod assay

On the morning after the week of running, running wheels were removed from the home cages and mice (wildtype) were tested on the accelerating rotarod. Mice were brought into a laboratory room designated for animal behavior and allowed to acclimatize for at least half an hour. Then, in groups of 4–5, mice were placed on a rotarod machine from Ugo Basile (model 47650), which contained a spinning rod that accelerated from 5 to 80 rotations per minute in 360 seconds. The latency to fall was recorded for each mouse as the time in seconds it took for the mouse either to fall off the rod or to grip onto the rod without walking for two full rotations. Trials were separated by 15 min. Mice were tested for nine trials on day 1, three trials on day 2, and three trials on day 9.

#### Slice electrophysiology

On the morning after the week of running, mice (DAT-cre/Ai9-tdTomato and ChAT-cre/Ai9-tdTomato) were anesthetized using isoflurane and perfused with ice-cold slicing solution that had been bubbled with 95% O2 and 5% CO2 (in nM: 198 glycerol, 2.5 KCl, 1.2 NaH2PO4, 20 HEPES, 25 NaHCO3, 10 glucose, 10 MgCl2, 0.5 CaCl2; 300–310 mOsm). Brains were extracted into oxygenated slicing solution and were sliced coronally at a thickness of 200 µm using a Leica vibratome (VT1000S). Slices containing SNc from the DAT-cre/Ai9 mice and containing DMS and caudal PPN from the ChAT-cre/Ai9 mice were incubated in holding solution (in nM: 92 NaCl, 2.5 KCl, 1.2 NaH2PO4, 30 NaHCO3, 20 HEPES, 2 MgCl2, 2 CaCl2, 35 glucose, 5 sodium ascorbate, 3 sodium pyruvate, 2 thiourea; 300–310 mOsm) at 34°C for 30 min then at room temperature for at least 30 min. After incubation, slices were perfused with artificial cerebrospinal fluid (ACSF; in nM: 125 NaCl, 25 NaHCO3, 3.5 KCl, 1.25 NaH2PO4; 300–310 mOsm) at 30–35°C in a recording chamber and visualized using an Evident OpenStand upright microscope. tdTomato fluorescence in neurons expressing cre allowed for identification of cholinergic neurons in ChAT-cre/Ai9 mice and dopaminergic neurons in DAT-cre/Ai9 mice using 565 nm ThorLabs LED light. Borosilicate glass micropipettes (World Precision Instruments, Inc. 1B150F-4, 4-inch length, 1.5 mm OD, 0.84 mm ID) of 1.5–4.5 MΩ resistance were created using a pipette puller (Sutter Instrument Model P-97) and were filled with a potassium methane sulfonate-based internal solution (in nM: 122 methanesulfonic acid, 9 NaCl, 9 HEPES, 1.8 MgCl2, 4 Mg-ATP, 0.3 Tris-GTP, 14 phosphocreatine; 285–295 mOsm). The internal solution also contained 0.3% neurobiotin to allow for post-hoc staining of recorded neurons. Whole-cell patch clamp recordings were performed in dopaminergic neurons of the SNc, CINs of the DMS, and cholinergic neurons of the caudal PPN. A Multiclamp 700B amplifier and Axon Digidata 1550B digitizer were used for electrophysiological record through the software Clampex 11.2. The signal was sampled at 10 kHz and was low-pass filtered at 2 kHz and 10 kHz for voltage clamp and current clamp, respectively.

In voltage clamp mode, cells were held at −70 mV for all recordings. A −5 mV step down to −75 mV was given for 100 ms to ensure proper break-in and to calculate input resistance and capacitance. To measure spontaneous excitatory postsynaptic currents (sEPSCs), 10-second recordings were obtained 20 times. Each sweep had a −5 mV step to ensure stable access throughout the protocol. The best 10 recordings were chosen for analysis based on the stability of the signal. sEPSC amplitude and frequency were analyzed post-hoc. Recordings in which access resistance exceeded 25 MΩ were excluded from analysis.

In current clamp mode, bridge was balanced and a gap-free recording with no current injection was performed to obtain the following measures: spontaneous action potential (AP) firing rate, coefficient of variation (CV) of spontaneous firing, average non-spike potential, average AP height, average AP width at half height, and average AP threshold. Then, a series of 4-second sweeps were collected in which a 1-second current injection was given starting 1 second into each sweep. The current injections ascended from −150 pA to +200 pA in increments of 50 pA, totaling eight sweeps. From these recordings, the following measures were obtained: average and maximum AP frequency at each depolarizing current injection, AP frequency gain (maximum frequency of current step ÷ average frequency during 0 pA step), adaptation index (1 - (frequency of last 4 APs of current step ÷ frequency of first 4 APs current step)), voltage sag (voltage at end of hyperpolarizing current step - voltage at beginning of hyperpolarizing current step), rebound delay (latency from end of hyperpolarizing current step until rebound firing), and rebound AP frequency. In SNc neurons, cells were considered in depolarization block if AP firing ceased within 0.5 seconds into the current step.

#### Immunohistochemistry and confocal imaging

After whole-cell recording, brain slices were fixed in 4% paraformaldehyde in 7.6 pH phosphate buffer (PB) and then cleared using the clear, unobstructed brain imaging cocktails (CUBIC) method as in^157,158^. At room temperature, slices were washed in PB, placed in CUBIC reagent 1 for one day, washed in PB, placed in blocking solution (0.5% fish gelatin in PB) for 3 hours, stained with streptavidin Cy5 for 3 days, washed in PB, and then placed in CUBIC reagent 2 for at least one hour before mounting and imaging.

A Leica SP8 laser confocal microscope at the Georgetown University Microscopy & Shared Resource core facility was used for imaging. Using LAS X software, z-stacks using a step size of 1µm were imaged of neurons stained with streptavidin. Cells that were too dim to reconstruct were excluded from analysis.

#### Cell reconstruction

Imaris image analysis software was used to perform full reconstructions of the three-dimensional neuron and dendritic arbor images. Axons were excluded from reconstruction. Cell reconstructions were done blinded to the exercise condition. Morphological characteristics of these reconstructions were extracted from Imaris and analyzed in Igor Pro. Sholl analysis^159^ calculated the number of dendritic intersections at each concentric radius from the soma in intervals of 1µm.

#### Estrous status determination

Following euthanasia of female mice for *ex-vivo* electrophysiology, vaginal lavage was performed using phosphate buffer solution. Cytology was examined under 10X magnification on an AmScope digital compound microscope to assess estrous status (Fig. S4A). As in^116^, proestrus was determined by a large proportion of nucleated epithelial cells. Estrus was determined by a large proportion of cornified epithelial cells. Metestrus was determined by relatively equal proportions of nucleated epithelial cells, cornified epithelial cells, and leukocytes. Diestrus was determined by a large proportion of leukocytes^160^.

### QUANTIFICATION AND STATISTICAL ANALYSIS

Custom scripts on Igor Pro were used to analyze electrophysiology data. The NeuroMatic package on Igor Pro was used to analyze spontaneous excitatory postsynaptic current data. Imaris and Igor Pro were used to analyze morphological data. Igor Pro was used to perform statistics and generate graphs. Two sample unpaired *t*-tests were used for statistical analysis. For comparisons where at least one group failed the Shapiro Wilk’s test for normality, Mann-Whitney-Wilcoxon *U*-tests were performed. One-way analysis of variance (ANOVA) was used for statistical analysis of comparisons with more than two groups. Fisher’s exact tests were used to determine differences between rates of depolarization block in SNc neurons. Pearson’s correlations were used to find the relationship between two variables across cells. For both depolarization block and correlation heat maps, the Benjamini-Hochberg procedure was used to control for false discovery rate with a Q value of 0.05. Fisher’s r-to-z transformations for independent correlations were used to calculate differences between correlations. For all data, samples reported are in the form of *N*=number of mice, *n*=number of cells.

## Acknowledgments

Funding for this research was provided by: National Institute on Aging (T32AG071745) grant awarded to V.J.L., BRAIN Initiative (R00NS112417) and Brain Research Foundation (BRFSG-2023-01) grants awarded to R.C.E., and Howard Hughes Medical Institute Gilliam Fellowship (GT1709) awarded to L.A.R. This research was supported by the Microscopy and Imaging Shared Resource (MISR, S10RR025661). We thank Dr. Thomas Coate for use of the Imaris reconstruction system and the Division of Comparative Medicine for their animal care and management.

## Author contributions

R.C.E. conceptualized the experiments; V.J.L. conducted electrophysiological experiments, tissue clearing and imaging, and analyzed data; L.A.R., G.S., and Z.A.C. conducted electrophysiological experiments; S.Q.Z., J.A.W., G.S., and A.F. performed tissue imaging and cell reconstructions; C.B.S. and O.M.K. performed behavioral experiments; M.R.C. and V.G.A performed pilot experiments; V.J.L. wrote the initial draft of the manuscript; V.J.L. and R.C.E. edited and revised the manuscript.

## Declaration of interests

The authors declare no competing interests.

## Notes

### Competing Interest Statement

The authors have declared no competing interest.

