## Supplemental Figures for "Sex-specific effects of exercise on motor coordination and extended basal ganglia physiology"

**Supplemental information titles and legends:**

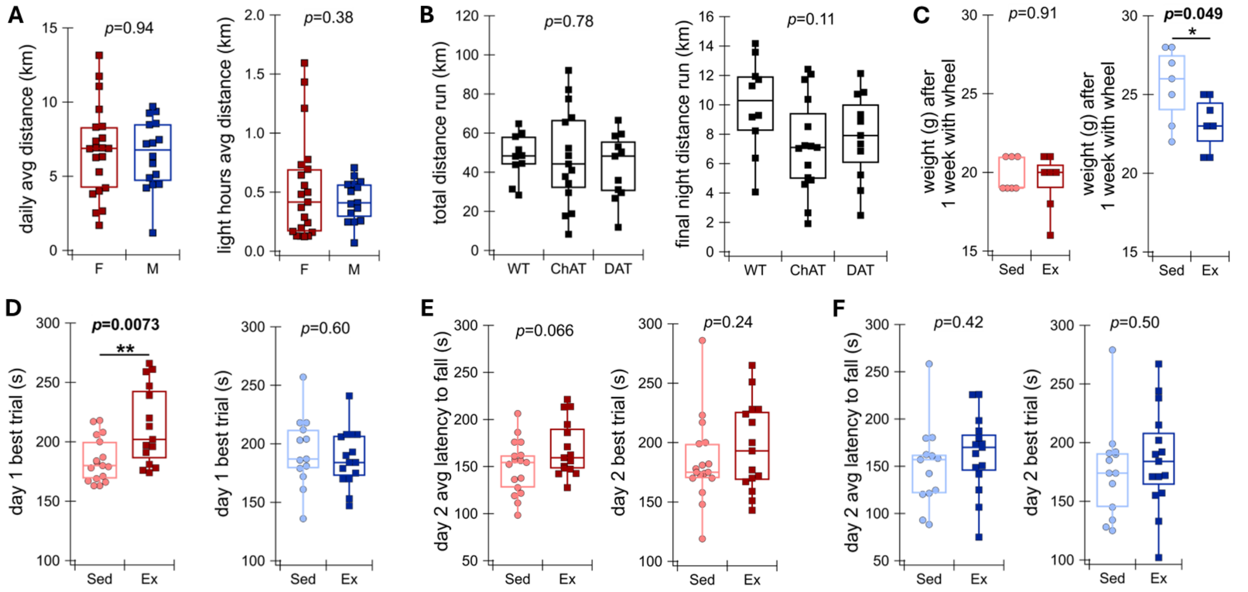


**Figure S1. Exercise enhancement of motor coordination is reduced by 24 hours after wheel removal, related to Figure 1.**

**(A)** Average daily distance run (left; *N*=21 Female (F): 6.73 ± 0.66 km, *N*=16 Male (M): 6.49 ± 0.59 km; Mann-Whitney-Wilcoxon *U*-test, *U*=165, *p*=0.94) and average light hours distance run (right; *N*=21 Female: 0.52 ± 0.094 km, *N*=16 Male: 0.43 ± 0.046 km; unpaired *t*-test, *t*=0.89, *p*=0.38) compared between males and females across this study.

**(B)** Total distance run across the week (left; *N*=10 Wildtype (WT): 47.92 ± 3.70 km, *N*=15 ChAT-cre/Ai9 (ChAT): 47.83 ± 6.45 km, *N*=11 DAT-cre/Ai9 (DAT): 42.83 ± 5.10 km, one-way ANOVA, *F*=0.25, *p*=0.78) and distance run on the final night (right; *N*=10 Wildtype: 9.98 ± 1.01 km, *N*=15 ChAT-cre/Ai9: 7.24 ± 0.85 km, *N*=11 DAT-cre/Ai9: 7.81 ± 0.90 km; one-way ANOVA, *F*=2.35, *p*=0.11) compared between all three genotypes in this study.

**(C)** Weight of wildtype female (left; *N*=7 Sedentary (Sed): 19.86 ± 0.40 g, *N*=7 Exercise (Ex): 19.43 ± 0.69 g; Mann-Whitney-Wilcoxon *U*-test, *U*=26, *p*=0.91) and male (right; *N*=7 Sedentary: 25.57 ± 0.90 g, *N*=7 Exercise: 23.14 ± 0.63 g; unpaired *t*-test, *t*=2.21, *p*=0.049) mice used in same rotarod experiment as Figure 1, as measured on day 1 of rotarod.

**(D)** Best trial of the 9 trials of day 1 in female (left; *N*=17 Sedentary: 184.65 ± 4.51 s, *N*=15 Exercise: 213.33 ± 8.56 s; unpaired *t*-test, *t*=-2.96, *p*=0.0073) and male (right; *N*=13 Sedentary: 193.38 ± 8.42 s, *N*=15 Exercise: 187.73 ± 6.25 s; unpaired *t*-test, *t*=0.54, *p*=0.60) groups in the same accelerating rotarod experiment as Figure 1.

**(E)** Average latency to fall (*N*=17 Sedentary: 149.39 ± 6.91 s, *N*=15 Exercise: 169.09 ± 7.66 s; unpaired *t*-test, *t*=-1.91, *p*=0.066) and best trial (*N*=17 Sedentary: 184.00 ± 8.68 s, *N*=15 Exercise: 196.33 ± 9.70 s; Mann-Whitney-Wilcoxon *U*-test, *U*=147, *p*=0.24) from the 3 trials on day 2 in females.

**(F)** Average latency to fall (*N*=13 Sedentary: 149.77 ± 12.24 s, *N*=15 Exercise: 163.00 ± 10.57 s; unpaired *t*-test, *t*=-0.82, *p*=0.42) and best trial (*N*=13 Sedentary: 175.46 ± 11.22 s, *N*=15 Exercise: 186.20 ± 11.15 s; unpaired *t*-test, *t*=-0.68, *p*=0.50) from the 3 trials on day 2 in males.

$*p < 0.05$, $**p < 0.01$.

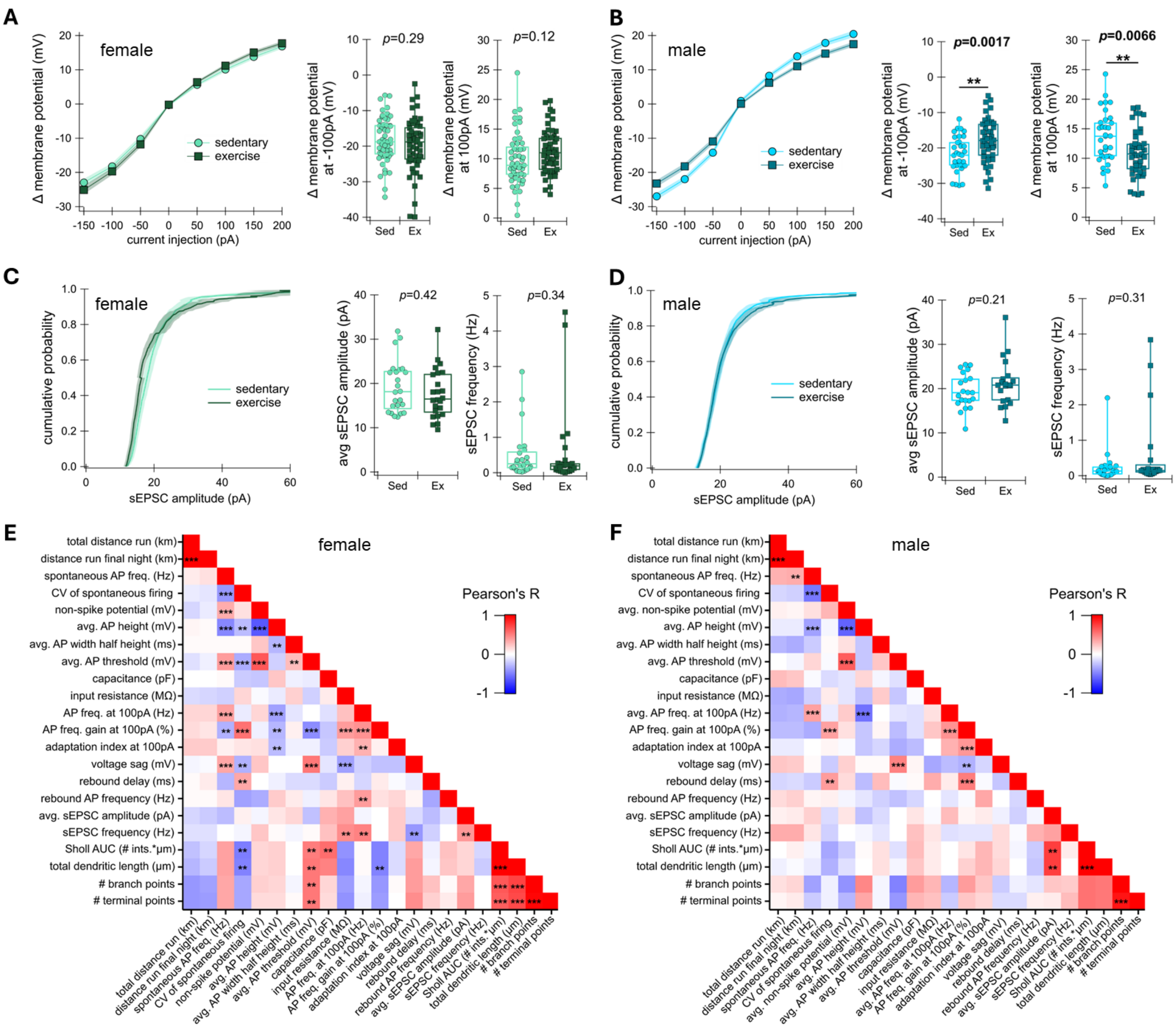


**Figure S2. Exercise reduces changes in membrane potential of SNc neurons selectively in male mice, related to Figure 2.**

**(A)** Left: Change in substantia nigra pars compacta (SNc) dopaminergic neuron membrane voltage from pre-injection voltage during current injections from -150pA to +200pA, increasing in increments of 50pA, compared between female groups in the same experiment as Figure 2. *n*=51 cells from 7 Sedentary mice (light green). *n*=55 cells from 7 Exercise mice (dark green). Middle: Change in membrane voltage during -100pA current injection in females (Sedentary (Sed): -18.27 ± 0.90 mV, Exercise (Ex): -19.71 ± 1.02 mV; unpaired *t*-test, *t*=1.053, *p*=0.29). Right: Change in membrane voltage during +100pA current injection in females (Sedentary: 9.96 ± 0.62 mV, Exercise: 11.19 ± 0.50 mV; unpaired *t*-test, *t*=-1.55, *p*=0.12).

**(B)** Same as in (A), but in males. Left: *n*=33 cells from 6 Sedentary mice (light blue). *n*=44 cells from 6 Exercise mice. Middle: Sedentary: -21.82 ± 0.87 mV, Exercise: -17.87 ± 0.93 mV; unpaired *t*-test, *t*=-3.11, *p*=0.0.0027. Right: Sedentary: 13.56 ± 0.76 mV, Exercise: 10.87 ± 0.59 mV; unpaired *t*-test, *t*=2.81, *p*=0.0066.

**(C)** Left: Cumulative probability of spontaneous excitatory postsynaptic current (sEPSC) amplitude in females. *n*=24 cells from 5 Sedentary mice (light green). *n*=24 cells from 6 Exercise mice (dark green). Middle: Average sEPSC amplitude in females (Sedentary: 19.20 ± 1.18 pA, Exercise: 17.66 ± 1.12 pA; Mann-Whitney-Wilcoxon *U*-test, *U*=328, *p*=0.42). Right: sEPSC frequency in females (Sedentary: 0.52 ± 0.14 Hz, Exercise: 0.58 ± 0.23 Hz; Mann-Whitney-Wilcoxon *U*-test, *U*=251, *p*=0.34).

**(D)** Same as in (C), but in males. $n$=21 cells from 6 Sedentary mice (light blue). $n$=20 cells from 5 Exercise mice (dark blue). Middle: Sedentary: 19.42 ± 0.86 pA, Exercise: 21.26 ± 1.18 pA; unpaired $t$-test, $t$=1.26, $p$=0.21. Right: Sedentary: 0.25 ± 0.098 Hz, Exercise: 0.61 ± 0.25 Hz; Mann-Whitney-Wilcoxon $U$-test, $U$=261, $p$=0.31.

**(E)** Heat map showing Pearson's correlations within each cell's electrophysiological and morphological properties as well as with each mouse's distance run in all (exercise and sedentary) females. Warmer colors indicate positive correlations and colder colors indicate negative correlations. The Benjamini-Hochberg procedure was used to control for false discovery rate.

**(F)** Same as in (E), but in males.

AP=action potential, AUC=area under the curve, avg=average, CV=coefficient of variation, freq=frequency, ints=Sholl intersections, sEPSC=spontaneous excitatory postsynaptic current

Data are represented as mean ± SEM. *$p < 0.05$, **$p < 0.01$, ***$p < 0.001$.

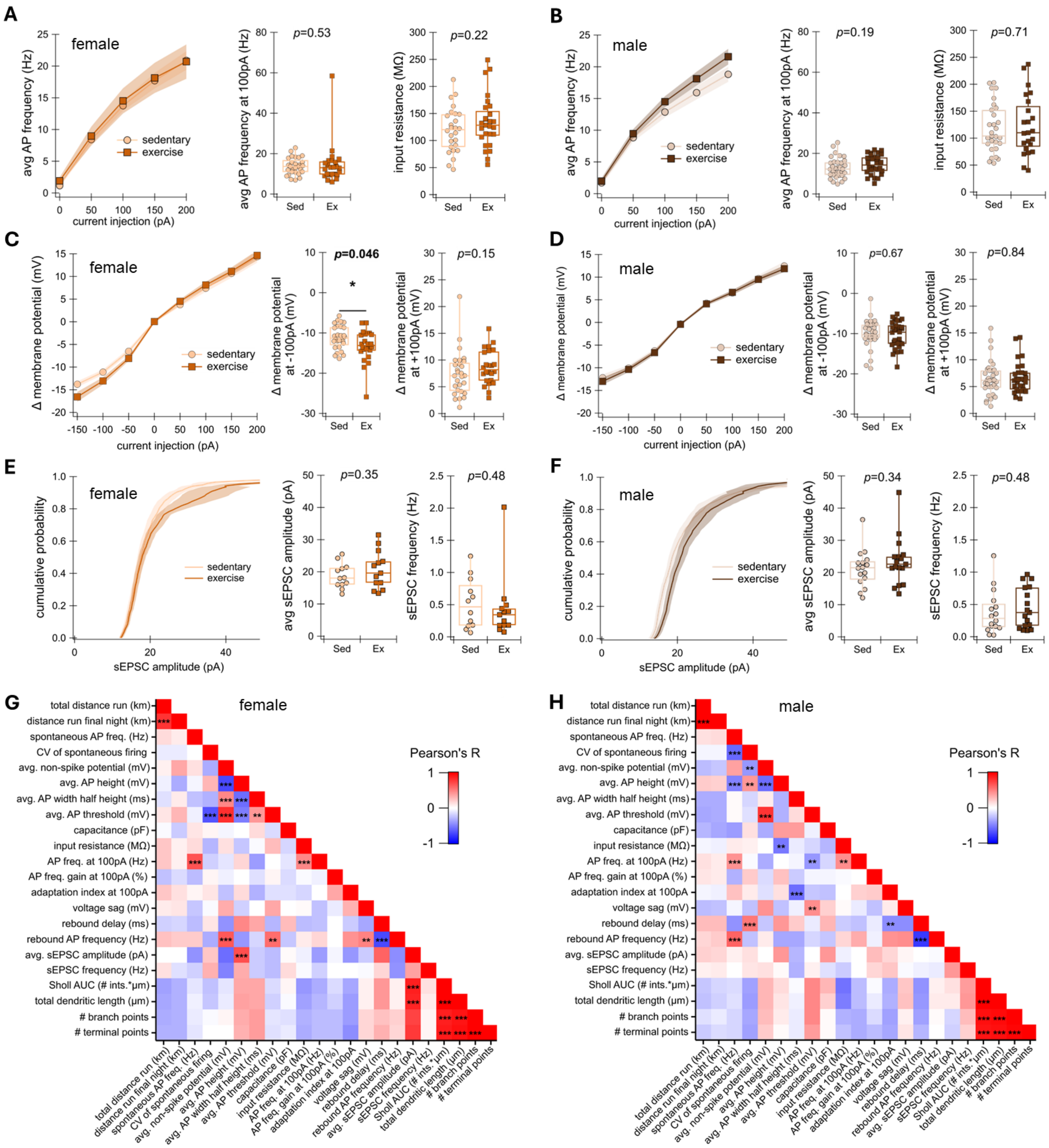


**Figure S3. Exercise has no effect on excitability or excitatory input measures of DMS CINs, related to Figure 4.**

**(A)** Left: Average action potential (AP) frequency of dorsomedial striatal (DMS) cholinergic interneurons (CINs) in female sedentary and exercise mice during each current injection step from 0 to 200, increasing in increments of 50pA, in the same experiment as Figure 4. *n*=28 cells from 8 Sedentary mice (light orange). *n*=24 cells from 8 Exercise mice (dark orange). Middle: Average AP frequency during the 100pA current step in females (*n*=28 cells from 8 Sedentary (Sed) mice: 13.75 ± 0.79 Hz, *n*=24 cells from 8 Exercise (Ex) mice: 14.51 ± 0.21 Hz; Mann-Whitney-Wilcoxon *U*-test, *U*=371, *p*=0.53). Right: Input resistance in females (*n*=26 cells from 7 Sedentary mice: 119.15 ± 8.09 MΩ, *n*=27 cells from 8 Exercise mice: 134.21 ± 9.26 MΩ; unpaired *t*-test, *t*=-1.23, *p*=0.22).

**(B)** Same as in (A), but in males. Left: $n$=33 cells from 8 Sedentary mice (light brown). $n$=30 cells from 7 Exercise mice (dark brown). Middle: $n$=33 cells from 8 Sedentary mice: 12.86 ± 0.86 Hz, $n$=30 cells from 7 Exercise mice: 14.49 ± 0.86 Hz; unpaired $t$=test, $t$=-1.34, $p$=0.19. Right: $n$=29 cells from 6 Sedentary mice: 117.89 ± 8.45 MΩ, $n$=23 cells from 5 Exercise mice: 123.16 ± 11.50 MΩ; unpaired $t$-test, $t$=-0.37, $p$=0.71.
**(C)** Left: Change in SNc neuron membrane voltage from pre-injection voltage during current injections from -150pA to +200pA, increasing in increments of 50pA, compared between female groups. $n$=28 cells from 8 Sedentary mice (light orange). $n$=24 cells from 8 Exercise mice. Middle: Change in membrane voltage during -100pA current injection in females (Sedentary: -11.12 ± 0.55 mV, Exercise: -13.28 ± 0.80 mV; Mann-Whitney-Wilcoxon $U$-test, $U$=227, $p$=0.046). Right: Change in membrane voltage during +100pA current injection in females (Sedentary: 7.40 ± 0.79 mV, Exercise: 8.65 ± 0.65 mV; Mann-Whitney-Wilcoxon $U$-test, $U$=257, $p$=0.15).
**(D)** Same as in (C) but in males. $n$=33 cells from 8 Sedentary mice (light brown). $n$=29 cells from 7 Exercise mice (dark brown). Middle: Sedentary: -10.07 ± 0.60 mV, Exercise: -10.37 ± 0.60 mV, Mann-Whitney-Wilcoxon $U$-test, $U$=527, $p$=0.67. Right: Sedentary: 6.50 ± 0.56 mV, Exercise: 6.64 ± 0.51 mV, Mann-Whitney-Wilcoxon $U$-test, $U$=480, $p$=0.84.
**(E)** Left: Cumulative probability of spontaneous excitatory postsynaptic current (sEPSC) amplitude in females. $n$=12 cells from 7 Sedentary mice (light orange). $n$=13 cells from 7 Exercise mice (dark orange). Middle: Average sEPSC amplitude in females (Sedentary: 18.75 ± 1.10 pA, Exercise: 20.59 ± 1.62 pA; unpaired $t$-test, $t$=-0.94, $p$=0.35). Right: sEPSC frequency in females (Sedentary: 0.52 ± 0.11 Hz, Exercise: 0.43 ± 0.14 Hz; Mann-Whitney-Wilcoxon $U$-test, $U$=91, $p$=0.48).
**(F)** Same as in (E), but in males. $n$=16 cells from 7 Sedentary mice (light brown). $n$=17 cells from 6 Exercise mice (dark brown). Middle: Sedentary: 21.19 ± 1.46 pA, Exercise: 23.40 ± 1.79 pA; Mann-Whitney-Wilcoxon $U$-test, $U$=109, $p$=0.34. Right: Sedentary: 0.38 ± 0.083 Hz, Exercise: 0.45 ± 0.076 Hz; Mann-Whitney-Wilcoxon $U$-test, $U$=116, $p$=0.48.
**(G)** Heat map showing Pearson's correlations within each cell's electrophysiological and morphological properties as well as with each mouse's distance run in all (exercise and sedentary) females. Warmer colors indicate positive correlations and colder colors indicate negative correlations. The Benjamini-Hochberg procedure was used to control for false discovery rate.
**(H)** Same as in (G), but in males.
AP=action potential, AUC=area under the curve, avg=average, CV=coefficient of variation, freq=frequency, ints=Sholl intersections, sEPSC=spontaneous excitatory postsynaptic current
Data are represented as mean ± SEM. *$p < 0.05$, **$p < 0.01$, ***$p < 0.001$.

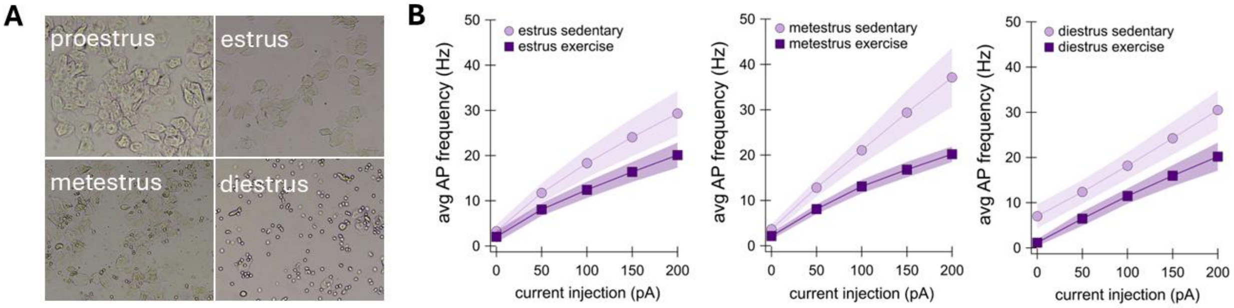


**Figure S4. Exercise effect on PPN cholinergic neuron excitability in females is not affected by estrous status, related to Figure 6.**
**(A)** Example images of vaginal cytology imaged at 10x showing the four estrous stages.
**(B)** Average AP frequency of caudal pedunculopontine nucleus (PPN) cholinergic neurons of females sedentary and exercise mice in estrus (left; $n$=9 cells from 3 Sedentary mice, $n$=6 cells from 1 Exercise

mouse), metestrus (middle; $n$=18 cells from 3 Sedentary mice, $n$=6 cells from 1 Exercise mouse), and diestrus (right; $n$=4 cells from 1 Sedentary mouse, $n$=6 cells from 2 Exercise mice) during each current injection step from 0 to 200, increasing in increments of 50pA, in the same experiment as Figure 6. Proestrus is not shown due to lack of samples.

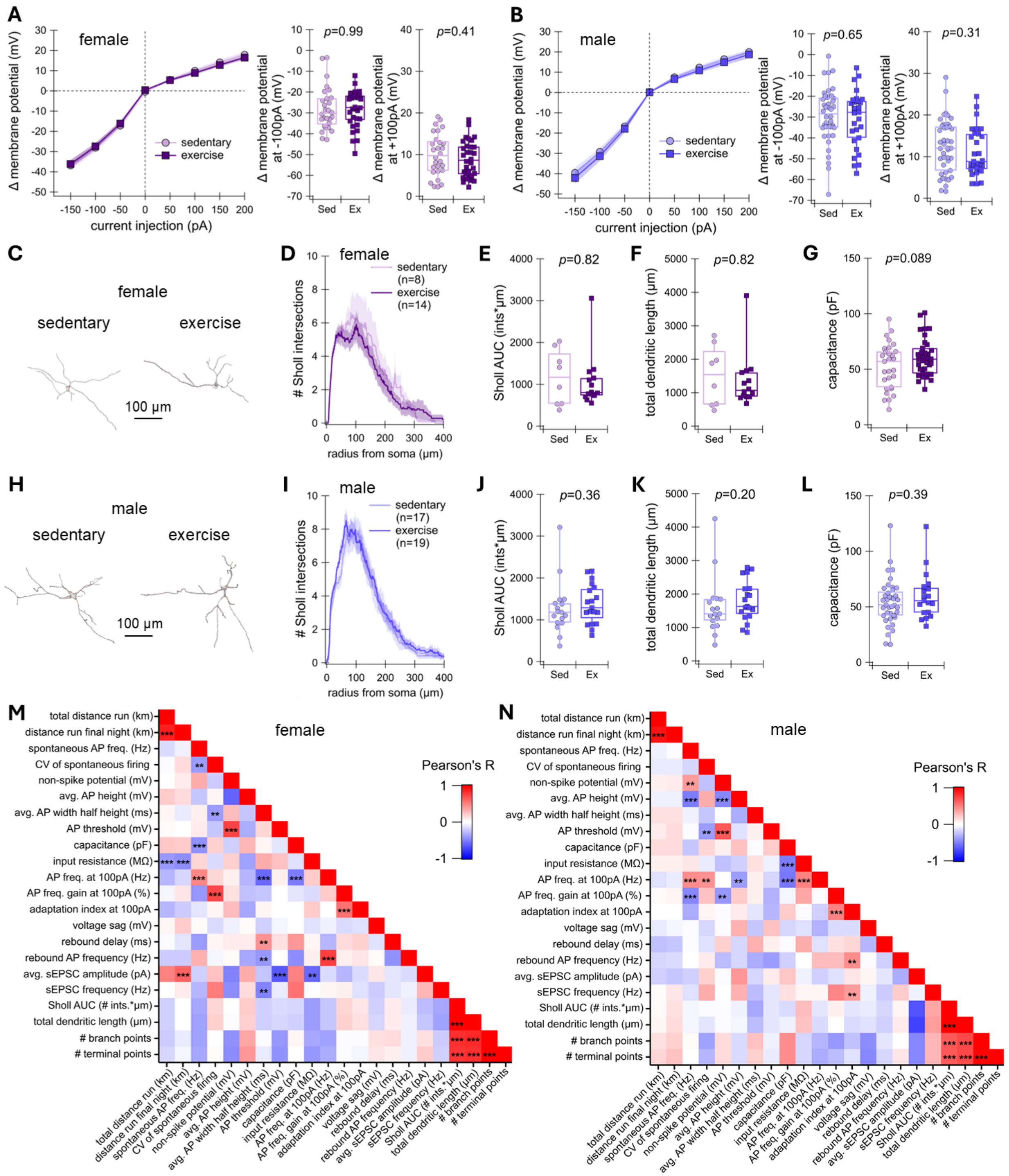


**Figure S5. Exercise has no effect on changes in membrane potential or morphology of PPN cholinergic neurons, related to Figure 6.**

**(A)** Left: Change in caudal pedunculopontine nucleus (PPN) cholinergic neuron membrane voltage from pre-injection voltage during current injections from -150pA to +200pA, increasing in increments of 50pA, compared between female groups. $n$=28 cells from 7 Sedentary mice (light pink). $n$=32 cells from 9 Exercise mice (dark pink). Middle: Change in membrane voltage during -100pA current injection in females (Sedentary (Sed): -28.11 ± 1.94 mV, Exercise (Ex): -28.11 ± 1.54 mV; unpaired $t$-test, $t$=-0.0019, $p$=0.99). Right: Change in membrane voltage during +100pA current injection in females (Sedentary: 9.94 ± 0.96 mV, Exercise: 8.93 ± 0.76 mV; unpaired $t$-test, $t$=0.83, $p$=0.41).

**(B)** Same as in (A) but in males. $n$=42 cells from 11 Sedentary mice (light purple). $n$=29 cells from 8 Exercise mice (dark purple). Middle: Sedentary: -29.38 ± 2.04 mV, Exercise: -30.74 ± 2.50 mV, unpaired $t$-test, $t$=0.45, $p$=0.65. Right: Sedentary: 12.39 ± 1.00 mV, Exercise: 11.06 ± 1.05 mV, Mann-Whitney-Wilcoxon $U$-test, $U$=696, $p$=0.31.

**(C)** Example reconstructed PPN cholinergic neurons in sedentary and exercise female groups.

**(D)** Number of Sholl intersections at each micron radius from the soma in sedentary (light pink; $n$=8 cells from 5 mice) and exercise (dark pink; $n$=14 cells from 5 mice) females.

**(E)** Area under the curve (AUC) for each cell's Sholl graph in females (Sedentary: 1162.37 ± 231.13 intersections*µm, Exercise: 1050.50 ± 168.01 intersections*µm; Mann-Whitney-Wilcoxon $U$-test, $U$=60, $p$=0.82).

**(F)** Total dendritic length in females (Sedentary: 1505.84 ± 312.65 µm, Exercise: 1343.74 ± 215.26 µm; Mann-Whitney-Wilcoxon $U$-test, $U$=60, $p$=0.82).

**(G)** Capacitance in females ($n$=29 cells from 6 Sedentary mice: 52.59 ± 3.85 pF, $n$=34 cells from 9 Exercise mice: 60.96 ± 2.92 pF; unpaired $t$-test, $t$=-1.73, $p$=0.089).

**(H)** Example reconstructed PPN cholinergic neurons in sedentary and exercise male groups.

**(I)** Number of Sholl intersections at each micron radius from the soma in sedentary (light purple; $n$=17 cells from 6 mice) and exercise (dark purple; $n$=19 cells from 7 mice) males.

**(J)** Area under the curve (AUC) for each cell's Sholl graph in males (Sedentary: 1272.65 ± 154.80 intersections*µm, Exercise: 1375.21 ± 107.24 intersections*µm; Mann-Whitney-Wilcoxon $U$-test, $U$=132, $p$=0.36).

**(K)** Total dendritic length in males (Sedentary: 1626.28 ± 211.03 µm, Exercise: 1801.99 ± 138.41 µm; Mann-Whitney-Wilcoxon $U$-test, $U$=203, $p$=0.20).

**(L)** Capacitance in males ($n$=39 cells from 9 Sedentary mice: 59.97 ± 3.27 pF, $n$=19 cells from 6 Exercise mice: 59.70 ± 4.83 pF; Mann-Whitney-Wilcoxon $U$-test, $U$=318, $p$=0.39).

**(M)** Heat map showing Pearson's correlations within each cell's electrophysiological and morphological properties as well as with each mouse's distance run in all (exercise and sedentary) females. Warmer colors indicate positive correlations and colder colors indicate negative correlations. The Benjamini-Hochberg procedure was used to control for false discovery rate.

**(N)** Same as in (M), but in males.

AP=action potential, AUC=area under the curve, avg=average, CV=coefficient of variation, freq=frequency, ints=Sholl intersections, sEPSC=spontaneous excitatory postsynaptic current

Data represented as mean ± SEM. *$p < 0.05$, **$p < 0.01$, ***$p < 0.001$.
